# Bridging Histology and Tractography: First In-Vivo Visualization of Short-Range Prefrontal Connections Informed by Primate Tract-Tracing

**DOI:** 10.1101/2025.10.22.683760

**Authors:** Matthew Amandola, Michael E. Kim, François Rheault, Bennett Landman, Kurt Schilling

## Abstract

Decades of histological research in non-human primates have revealed a dense web of short-range connections underpinning prefrontal cortex (PFC) function. However, translating this anatomical ground-truth to the living human brain has been a major challenge, leaving our understanding of the PFC’s intrinsic wiring incomplete. These short-range fibers are difficult to resolve with non-invasive methods like diffusion tractography, which are often hampered by false positives. Here, we provide the first systematic in-vivo visualization of these pathways in the human brain. By informing high-resolution probabilistic tractography with established tract-tracing findings, we mapped 91 histologically-defined short-range connections within and between five major PFC subdivisions in 1,003 individuals (547 F, 456 M). Our anatomically-informed approach successfully reconstructed these intricate connections with high precision (>80%) and accuracy (>70%) relative to histological findings. The resulting tracts not only captured broad organizational principles but also replicated fine-grained patterns previously only seen in invasive studies. Furthermore, these connections showed high test-retest reliability within individuals alongside significant variability between them, highlighting a stable yet unique anatomical fingerprint. Ultimately, this study shows how linking histology to tractography provides a powerful framework to advance our understanding of the human connectome and opens avenues to investigate local circuitry that underpins cognition and disease.

## 1. Introduction

The prefrontal cortex (PFC) of the human brain consists of a number of different cortical regions anterior to the precentral gyrus grouped by their organizational properties and cytoarchitecture (Brodmann, 1909; Petrides et al., 2012). The PFC is the most advanced association region of the human brain and is crucial for complex cognitive functioning, including working memory (Levy & Goldman-Rakic, 2000) and executive functioning (Funahashi et al., 1993). To facilitate these cognitive tasks, the PFC is highly structurally and functionally connected to other association regions of the cortex, such as the posterior parietal cortex and motor association areas (Petrides & Pandya, 1989; Marek & Dosenbach, 2018). These connections typically come in the form of long-range, deep white-matter association pathways (Schmahmann & Pandya, 2006).

In addition to long-range connections, the PFC displays a high degree of interconnectivity via smaller, short-range connections called short association fibers (SAFs). These shorter-range connections connect adjacent gyri, sulci, and different cytoarchitectural delineations within the PFC, allowing these regions to communicate, as well as supporting the local processing necessary for integrating information and guiding flexible behavior (Haber et al., 2022). However, the intrinsic connectivity of the PFC remains unclear in the human brain, including the patterns of fine-scale organization of short-range connections, their particular cortical terminations, and their role in supporting information transfer underlying cognition and behavior. While prior work has approximated broad connectomic trends across PFC subregions, precise knowledge of which cytoarchitecturally defined regions are structurally interconnected in the human brain remains incomplete. Conventionally, the short-connections of the PFC have been studied using tract-tracing in non-human primates (Barbas & Pandya, 1989; Haber et al., 2022). These tract-tracing studies suggest clear, replicable patterns of structural connectivity between the differing cytoarchitectural divisions of the PFC (Barbas & Pandya, 1989; Yeterian et al., 2012; Haber et al. 2022). Additionally, a selection of these short-range PFC connections are replicated in the human brain using post-mortem Klinger dissection (Catani et al., 2012), suggesting the organization of these intrinsic connections are similar across species. While tract-tracing and dissection studies have been illuminating in the mapping of the intrinsic connections of the PFC, there are considerable limitations to these methods. Tract-tracing literature, while precise, is limited by the fact that histological findings are cross-species, making the extent that we can generalize results to the human brain unclear. Further, dissection lacks the precision to resolve fine-grained pathways. Taken together, this points to a need for a precise, non-invasive, in-vivo method to study the intrinsic connections of the PFC in the human brain.

Currently, the only way to measure white-matter tracts of the human brain in-vivo is with diffusion MRI fiber tractography. However, tractography is prone to generating a high number of false positive connections (Maier-Hein et al., 2017), a challenge compounded by the lack of an established ground truth for human structural connectivity (Dyrby et al., 2007). Encouragingly, previous tractography studies have succeeded in mapping accurate long-range connections in the human brain when modelling connections between regions informed by evidence from tract-tracing and dissection (De Schotten et al., 2011; Girard et al., 2020; Schilling et al., 2020; Amandola et al., 2025). Further, while there are fewer tractography studies examining SAF’s, these studies show promising reconstruction of these shorter range pathways (Guevara et al., 2022; Pietrasik et al., 2023; Schilling et al., 2023; Van Dyken et al., 2024), especially when cross-validating these SAF’s by tract-tracing or dissection (Catani et al., 2012). This suggests that tractography informed by the histological literature can become a powerful tool in observing these under-investigated pathways in the human brain.

Here, we leverage the rich primate histological literature as a biological blueprint to inform high-resolution tractography in a large-scale cohort, providing the first systematic in-vivo visualization of the PFC’s short-connections. While this study does not directly utilize NHP tract-tracing data, this extensive body of work informed our region of interest (ROI) selection, defined our morphological expectations for bundle reconstruction, and provided a benchmark for evaluating the anatomical fidelity of the results. The goal of the current study is to elucidate the patterns of structural interconnection within the PFC, with the explicit goal of mirroring the precision of the many seminal works from the tract-tracing literature. Here, we aim to clearly delineate which short-range pathways can be reliably reconstructed in-vivo, with the long-term objective of establishing a canonical set of prefrontal SAFs analogous in clarity and anatomical definition to the well-characterized long-range white-matter tracts of the brain. Our approach successfully reconstructs these intricate pathways, modeling both the presence and principled absence of connections with high fidelity to the histological findings. We show that these connections are highly reproducible within individuals yet exhibit variability across the population, revealing a stable and individually unique anatomical fingerprint. In sum, this work provides a framework for mapping a previously inaccessible level of white-matter architecture, bridging findings from animal models to human neuroscience, and advancing understanding of how local cortical wiring supports cognition and disease.

## 2. Methods

### 2.1 Subjects

Brain images from 1065 subjects were downloaded from the Human Connectome Project Young Adult (HCP) Study (van Essen et al., 2012) in October 2024. A total of 1003 subjects were included in the study (547 F, 456 M; age range = 22 - 36). HCP Young Adult is one of the datasets available from the Human Connectome Project (HCP), which is a multi-site, large-scale neuroimaging project which collects functional, structural, and diffusion magnetic resonance imaging (MRI) data, as well as physiological and behavioral data (Bookheimer et al., 2019). Inclusion criteria for our study was consistent with the criteria listed from HCP: No significant history of psychiatric disorder, substance abuse, neurological, or cardiovascular disease. No report of diagnosis by a treating physician. No pharmacologic or behavioral treatment for 12 months or longer by a specialty-trained physician (psychiatrist, neurologist, cardiologist) or therapist (e.g., psychologist, social worker). Subjects must have the ability to give valid informed consent, and have normal cognitive abilities.

### 2.2 Imaging Parameters

Both structural T1 and diffusion MR contrast were acquired with a Siemens 3T Prisma whole-body scanner (32-channel head coil). MR images for this HCP sample were collected from four different scan sites. T1 images had 256 sagittal slices with slice thickness = .7 mm, were acquired using a single-echo MPRAGE sequence, with repetition time (TR) = 2400 ms, inversion time (TI) = 1000 ms, echo time (TE) = 2.14 ms, voxel size = .7 × .7 × .7 mm^3^, flip angle = 8°, matrix = 320 × 320, field of view (FOV) = 224 × 224.

We used the minimally preprocessed data (Glasser et al. 2013) from HCP Q1-Q4 2015 release, which uses FSL TOPUP and EDDY algorithms to conduct susceptibility correction, motion correction, and eddy current corrections. Multishell Diffusion MR sequences were collected with multi-band = 3, b = 1000, 2000, and 3000, 90 directions per shell, TR = 5520 ms, TE = 89.5 ms, voxel size = 1.25 × 1.25 × 1.25 mm^3^, flip angle = 78°, matrix = 168 × 144, FOV = 210 × 180 mm.

### 2.3 Cortical Parcellation

For ROI selection, we used the Human Connectome Project Multi-Modal Parcellation 1.0 atlas (Glasser et al., 2016). These ROI’s were transformed from surface space to Freesurfer’s fsaverage space (Fischl, 2012), and then from each subject’s fsaverage space to their volumetric space. This resulted in precise, individualized Brodmann ROI’s for each subject. We then followed the five canonical partitions of the prefrontal cortex (for review, see Haber et al., 2022): the dorsolateral PFC (dl-PFC), which consisted of Brodmann Areas 8, 9, 9/46, and 46; the ventrolateral PFC (vl-PFC), which consisted of Brodmann Areas 44, 45, and 47; the frontal pole, which consisted of Brodmann Area 10; the orbitofrontal cortex (OFC), which consisted of Brodmann Areas 11, 13, and 14; and the anterior cingulate cortex (ACC), which consisted of Brodmann Areas 24, 25, and 32. For a full overview of the subregions each ROI consists of, see Table 1. The specific anatomical connections reconstructed for each PFC partition are detailed in the *Anatomical Results and Test-Retest* section.

**Table 1:** Definitions of PFC Regions using the HCP-MMP1 Atlas.

| **PFC Region** | **Area** | **Constituent HCP-MMP1 ROI's** |
| --- | --- | --- |
| **dl-PFC** |  |  |
|  | 8 | 8Ad, 8BM, 8BL, 8C |
|  | 9 | 9m, 9a, 9p |
|  | 46 | 46 |
|  | 9/46 | 9-46d, 9-46av, 9-46pv |
| **vl-PFC** |  |  |
|  | 44 | 44 |
|  | 45 | 45 |
|  | 47 | a47r, p47r, 47s, 47l, 47m |
| **OFC** |  |  |
|  | 11 | 11l |
|  | 13 | 13l |
|  | 14* | OFC, pOFC |
| **Frontal Pole** |  |  |
|  | 10 | 10r, 10d, 10v, ,10pp, a10p, p10p |
| **ACC** |  |  |
|  | 24 | 24a, 24p, a24pr, p24pr, 24dd, 24dv |
|  | 25 | 25 |
|  | 32 | 32d, 32p, 32s, a32pr, p32pr |

\* = Because the HCP-MMP1 atlas does not contain parcels exclusive to area 14, we used the OFC and pOFC parcels, which encompass portions of areas 11 and 13. Since areas 11 and 13 have dedicated parcels elsewhere in the atlas, we felt this was the most suitable way to approximate area 14.

### 2.4 Tractography Parameters

For each subject, anatomically-constrained tractography was performed using the MRTrix3 software (Tournier et al., 2019), using the iFOD2 algorithm. All diffusion data were resampled to 1mm^3^, and we conducted multi-shell, multi-tissue constrained spherical deconvolution using MRTrix3’s dwi2fod command (Jeurissen et al., 2014). We then used MRTrix’s 5TTGen command to create a five tissue type (5TT) image from each subject’s structural image, which allowed us to extract the gray matter white matter interface (GMWMI) volume for every subject. Tractography was performed using the second-order integration probabilistic algorithm (Tournier et al., 2019), with default parameters to generate 1,000 streamlines per run, though max length varied from 38-125mm per tract based on preliminary testing, histological findings, and anatomical boundaries in order to ensure the highest probability of short-range fiber selection, rather than deep white matter. We conducted preliminary testing and thorough manual assessment before deciding on final maximum lengths. Using a sub-sample of 10 subjects, we conducted PFC tractography without implementing a maximum length. We then visualized the reconstructed tracts for each subject in the MI-Brain GUI (Rheault et al., 2016) and evaluated if (i) each individual modeled tract reached both Brodmann ROIs, (ii) the tract did not include streamlines branching from any deep white matter structure, and (iii) the tract was not dominated by false positive streamlines. If the reconstructed tracts picked up streamlines from nearby deep white matter structures or had an excess of false positives, we then manually shortened the maximum length for the given tract using the MI-Brain GUI until the tracts satisfied all three requirements for each individual in the testing cohort. Lastly, to choose final maximum lengths for the entire cohort, we averaged the empirically determined maximum length across the 10 individuals for each tracts plus 1.5 standard deviations to account for individual variability. For a full overview of the final chosen maximum lengths, see Supplementary Table 1. Tractography was run between each ROI pair between the dl-PFC, vl-PFC, frontal pole, OFC, and ACC, as well as between ROI pairs within each PFC partition. This resulted in a total of 91 tracts for each subject. We chose our streamline count based on precedence set by previous literature. TractSeg, one of the most widely-employed tractography bundle reconstruction methods, employs a default streamline count of 2,000 for large, complex, deep-white matter tracts (Wasserthal et al., 2018). Given that the SAFs examined here are relatively simple, local trajectories, we decided using this established streamline count would be reasonable to capture these pathways while limiting the generation of spurious streamlines. For each tract we masked the subject’s GMWMI volume by the first ROI in the corresponding ROI pair, and seeded from the masked GMWMI image with the second ROI as the target with a maximum streamline count of 1,000. We then repeated this procedure with ROI’s flipped, and summed the two tracts, for a maximum final streamline count of 2,000. For example, for the tracts between BA 9 and BA 46, we would mask the GMWMI volume by BA 9, and seed from this masked GMWMI with BA 46 as a target. Then, we would mask the GMWMI volume by BA 46, seed from this masked GMWMI with BA 9 as a target. Finally, we concatenated the two tracts and performed outlier removal using scilpy’s *scil_bundle_reject_outliers* function (Renauld et al., 2026) on the resulting tract, and each tract was then visually inspected.

### 2.5 Comparison to Histology

To evaluate our tractography results against the primate histological literature, we classified each potential pathway based on its anatomical plausibility, reproducibility, and concurrence with histological literature. Histology has two classifications:

- **Present in histology.** This categorization represents a consistent representation of a connection between two areas throughout histological literature, and connections which have been corroborated in multiple studies. It is important to note, however, that despite this classification there is no complete ground truth in literature. These connections have generally been accepted as existing between individuals.
- **Absent in histology.** This categorization represents connections that do not have any histological precedence. This can also include a limited number of connections that are reported in only singular instances, inconsistent, or are a result of a spread of the tracer in the injection site to adjacent cortical areas.

Out of a possible 91 studied connections, histological literature suggests 72 true positive connections and 19 true negative connections (Pandya et al., 1981; Barbas & Pandya, 1989; Preuss & Goldman-Rakic, 1989; Carmichael & Price 1995a; Carmichael & Price 1995b; Carmichael & Price 1996; Petrides & Pandya 1999; Cavada et al., 2000; Semendeferi et al., 2001; Petrides & Pandya, 2002; Petrides & Pandya, 2006; Schmahmann & Pandya 2006; Petrides & Pandya, 2007; Gerbella et al., 2010; Morecraft et al., 2012; Yeterian et al., 2012; Frey et al., 2014; Haber et al., 2022). For a full overview, see Figure 1.

**Figure 1.**
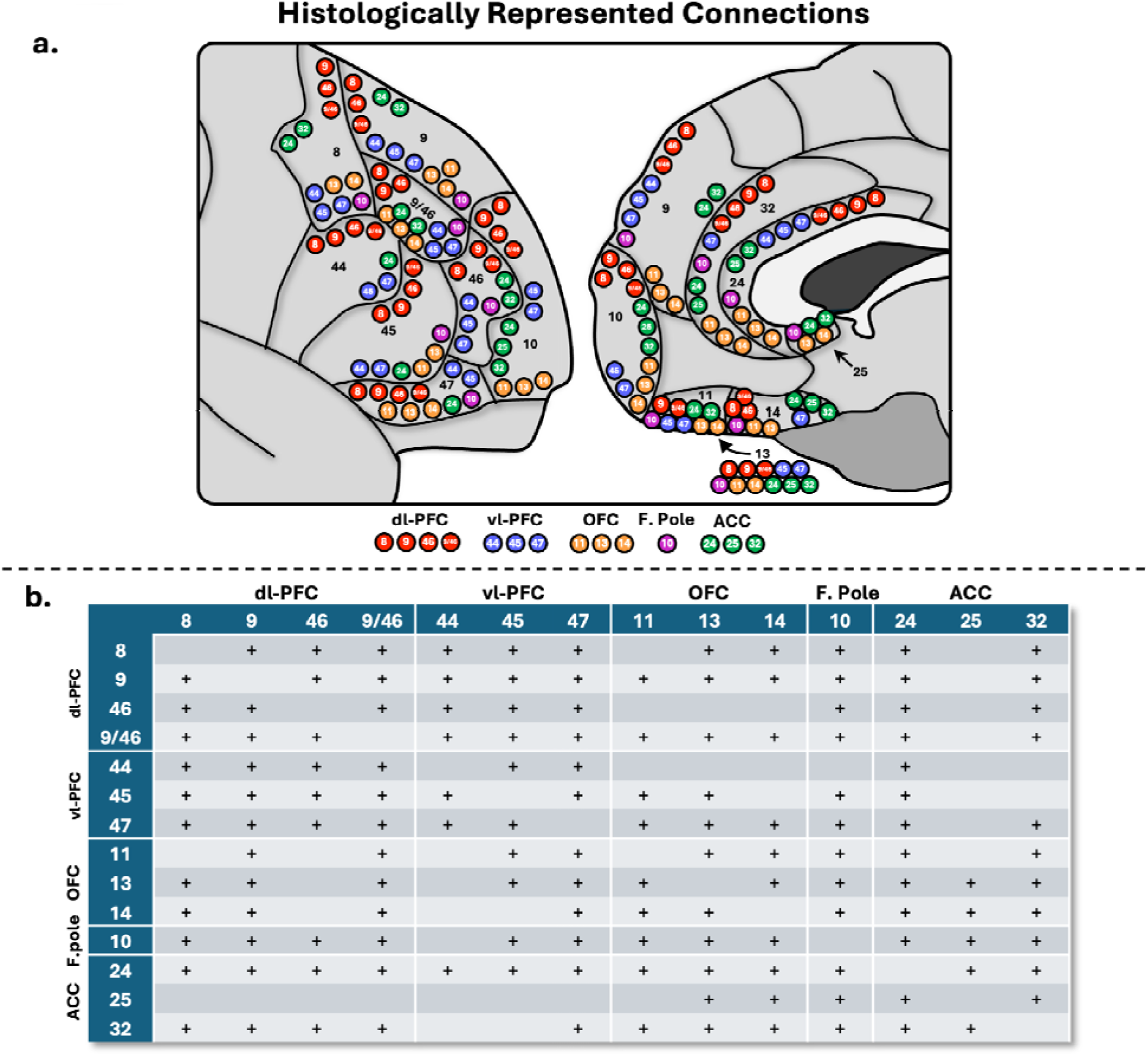
a.) Schematic depicting the interconnections of the prefrontal cortex (PFC). Each dot represents a connection with histological precedence. Red = dl-PFC (dorsolateral prefrontal), blue = vl-PFC (ventrolateral prefrontal), orange = orbitofrontal, purple = F. Pole (frontal pole), green = ACC (anterior cingulate). b.) Table overview of histologically supported connections. + = consistently shown in histological literature.

For our tractography classifications, we performed a visual QA and assessed anatomical plausibility and consistency across the population. Based on this, we classified individual tractography bundles into 3 classifications:

- **Anatomically plausible bundles.** Bundles that followed coherent trajectories aligned with anatomical landmarks, demonstrated smooth directional continuity, and displayed few branching streamlines. Tracts were classified as anatomically plausible only if they satisfied all of the following: i.) High relative density of streamlines; ii.) High relative number of streamlines reconstructed; and iii.) Adherence of the reconstructed bundle to local anatomy. These criteria were informed by prior work using similar QA frameworks (Vavassori et al., 2025) and based on our combined knowledge and experience with neuroanatomy and diffusion image processing.
- **Anatomically implausible bundles.** Tracts classified as anatomically implausible failed to meet one or more of the above criteria and typically exhibited well-described features of false positive tractography, including abrupt directional changes, incongruity with surrounding anatomy, random trajectories, or streamline looping.
- **Absent in tractography.** This categorization represents bundles that produced no or few streamlines.

To ensure a conservative and accurate representation of tractography performance, each hemisphere was inspected independently for classification. This approach was chosen to maintain methodological parsimony and interpretability of results, as comparisons revealed that reconstruction success was highly symmetrical across the population.

Based on these classifications of anatomically plausible, anatomically implausible, and absent tracts for each subject, we assigned bundles to one of four categories as detailed below and in Figure 2.

- **True Positive (TP).** Bundles which have histological precedence, are reconstructed in greater than 50% of the subject pool, and are anatomically plausible in tractography. In order to gauge how consistent across subjects these true positive bundles were, we created a robust sub-category:

- **Robust True Positive (RTP)**: Pathways with established histological precedent that were extremely consistent across individuals, present in over 80% of subjects.
- **True Negative (TN).** Bundles which do not have histological precedence and are absent in tractography.
- **False Positive (FP).** Bundles which do not have histological precedence, but are reconstructed in over 50% of the subject pool (anatomically plausible and anatomically implausible trajectories) Note, this also includes false positive trajectories where there is *histological precedence*, but tractography follows a clearly anatomically implausible trajectory^1^ in over 50% of the subject pool (Maier-Hein et al., 2017).
- **False Negative (FN).** Bundles which have histological precedence but are absent in tractography.

**Figure 2.**
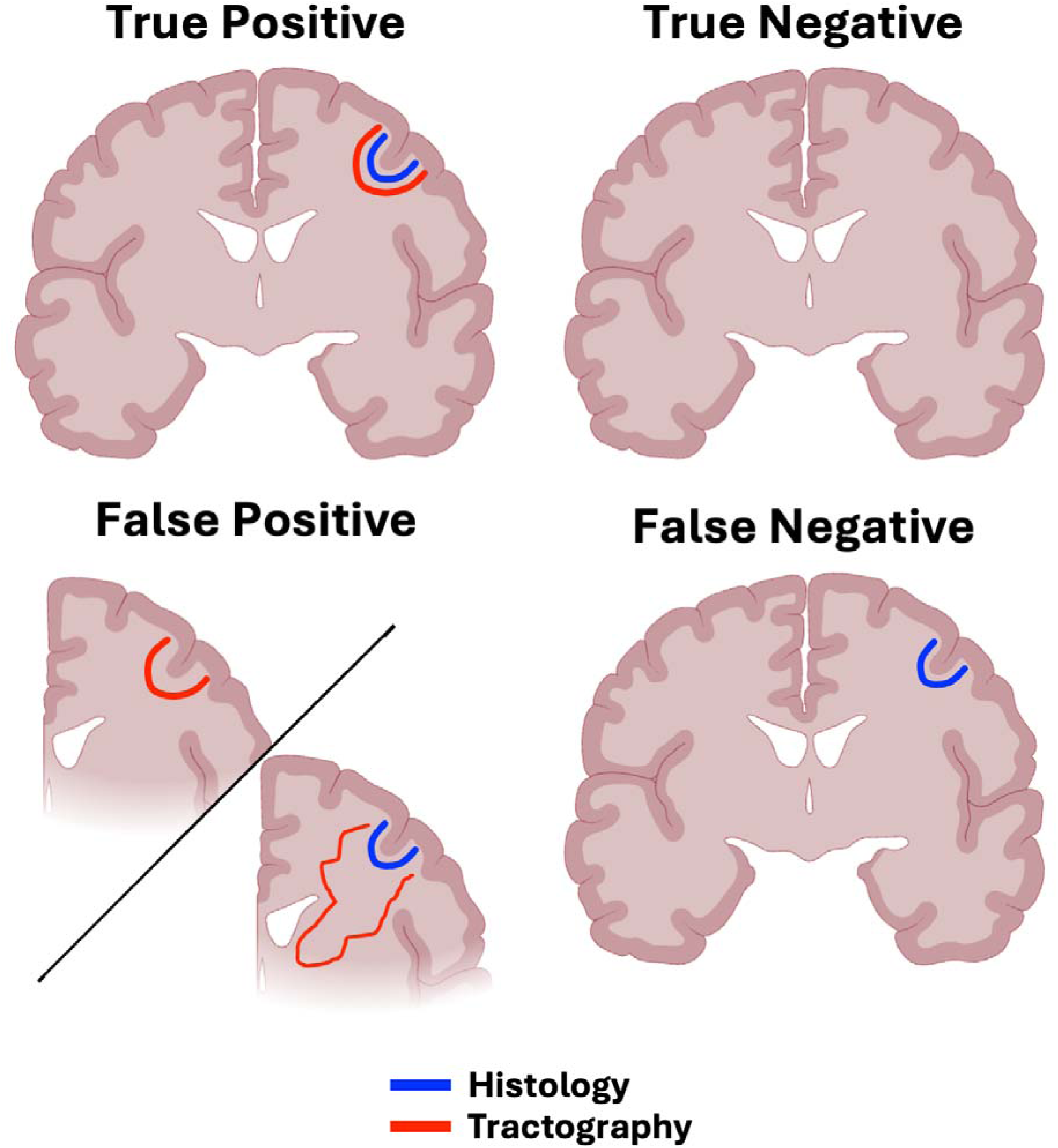
Classifications of tractography outputs in comparison to histology. Blue = connection with histological evidence, red = tractography trajectories of the same connection. True positive = has histological precedence and tractography bundles are both anatomically plausible and reconstructed in over 50% of the subject pool; true negative = no histological precedence and absent in tractography; false positive = no precedence in histology but reconstructed in tractography, or anatomically implausible tractography trajectory consistent across subjects regardless of histological precedence; false negative = has histological precedence but absent in tractography.

For the purpose of calculating summary statistics, TP and TN results were considered to be in agreement with the histological literature, while FN and FP results were considered to be in disagreement. To review, each individual tract is visually inspected to determine if it meets criteria for anatomically plausible, implausible, or absent for each subject. Then, once this tract has been visually inspected across the whole subject pool, it is assigned to one of the classifications above. For examples of each classification, see Supplementary Figures 1-5.

### 2.5 Test-retest Reliability

To assess the reproducibility of our findings, we quantified two distinct forms of variability:

- **Within-subject reliability** was evaluated using the HCP test-retest dataset (*n* = 44 subjects; 31 F, 13 M, age range 22-35). For these subjects, the entire tractography pipeline was run independently on two separate diffusion scans to measure the consistency of the reconstructed pathways within the same individual.
- **Between-subject variability** was quantified by comparing the same reconstructed pathways across different individuals aligned in MNI space (Fonov et al., 2011). We implemented ANTS antsRegistrationSyN to create nonlinear deformation files (Avants et al., 2008), coregistering each subject’s T1 image to MNI space. We then applied this nonlinear transformation directly to the streamlines using *scil_tractogram_apply_transform* from scilpy (Renauld et al., 2026). This way, we avoid common distortion problems associated with track-density map transformations.

For both analyses, we used two metrics to compare the spatial characteristics of the reconstructed bundles using the *scil_bundle_pairwise_comparison* function in scilpy (Renauld et al., 2026):

- **Weighted Dice (wDice)**: wDice is a measure of volumetric overlap, particularly sensitive to the spatial correspondence of the high-density core of the tracts. In comparison to the traditional Dice coefficient (Dice, 1945), wDice was specifically designed for fiber tracts measured by diffusion imaging by incorporating both spatial proximity and streamline density information. Rather than treating voxels as binary, wDice weights them according to the local density of streamlines and their spatial correspondence. This makes wDice a more sensitive and anatomically meaningful measure of reproducibility for small or spatially variable fiber pathways (Cousineau et al., 2017; Zhang et al., 2019a; Rheault et al., 2020). wDice scores range from 0 (no overlap) to 1 (perfect overlap) (Cousineau et al., 2017; Zhang et al., 2019a; Rheault et al., 2020).
- **Bundle Adjacency:** A measure of spatial disagreement, calculated as the average distance between the non-overlapping portions of two bundles, similar to Hausdorff distance (Schilling et al., 2021a). A lower value indicates better geometric alignment; for example, a value of 3mm means that where the bundles differ, their volumes are, on average, 3mm apart.

## 3. Results

### 3.1 Correspondence with Histological Findings

To quantify the performance of our histology-informed tractography, we evaluated each reconstructed pathway against the primate literature using the classification scheme defined in the Methods. From these classifications (TP, TN, FN, FP), we calculated the overall accuracy, sensitivity, specificity, and precision of the reconstructions across the entire PFC and within its major subdivisions.

Overall, our tractography results showed strong correspondence with the histological literature. Throughout the whole PFC (91 potential pathways), 49 total tracts were deemed true positives (41 RTP), 17 tracts were deemed true negatives, against 12 tracts false negatives and 13 false positives. This yielded an overall high accuracy of 73% and a sensitivity of 80% for detecting established connections. Notably, the method demonstrated strong precision (79%) and moderate specificity (57%), indicating a strong robustness to false-positive tracts.

Performance varied across the PFC subdivisions: The tracts of the dl-PFC displayed similar results to the entire PFC. The tracts of the dl-PFC resembled histological findings, as after visual inspection, the regions from the dl-PFC showed 26 true positive tracts (22 RTP), 8 true negative tracts, 6 false positive tracts, and 6 false negative tracts, for an overall 74% total accuracy. The dl-PFC also had 82% sensitivity and 82% precision, however, specificity was the lowest at 57%.

The tracts of the vl-PFC also performed well: the vl-PFC displayed 20 true positives (17 RTP), 7 true negatives, 6 false positives, and 3 false negatives, for an overall 75% accuracy, 87% sensitivity, 77% precision, and 54% specificity with respect to histological findings.

After visual inspection, the regions from the OFC had 14 true positives (10 RTP), 8 true negatives, 4 false positives, and 10 false negatives, for an overall 61% accuracy with respect to histological literature. This is driven by a low sensitivity (58%), as the OFC had the most false negatives out of all regions. Yet, the OFC still displayed moderate specificity (66%), and high precision (78%).

The connections of the frontal pole showed perfect histological correspondence, with 12 true positives (9 RTP), 1 true negative, and no false positives or negatives, resulting in 100% accuracy, sensitivity, specificity, and precision.

After visual inspection, the regions from the ACC had 13 true positives (9 RTP), 8 true negatives, 9 false positives, and 6 false negatives (see Figure 7), with an overall 58% accuracy with histological literature. This makes the ACC the region with the lowest accuracy, despite its relatively high sensitivity (68%). The ACC was also the most susceptible to false positive tracts, with a specificity of 47%, and a precision of 59%.

In summary, histology-informed tractography showed high accuracy, sensitivity, and precision, but moderate specificity, when reconstructing the short range fibers of the PFC. The dl-PFC, vl-PFC, and especially the FP, closely resembled histological findings, displaying high accuracy. The OFC and ACC still performed relatively well, but were sensitive to false negatives and false positives, respectively.

### 3.2 Anatomical Results and Test-Retest

We now present the detailed anatomical and reproducibility findings, organized by the five major PFC partitions. For each prefrontal region (Figures 3-7), the results are presented in a consistent structure. We first provide a qualitative description of the general organization of the reconstructed pathways for the entire subdivision. We then provide a detailed, area-by-area account of the specific cortico-cortical connections. Finally, this is followed by the quantitative results of our reproducibility analyses, reporting the within-subject (test-retest) reliability and between-subject variability for the tracts.

**Figure 3.**
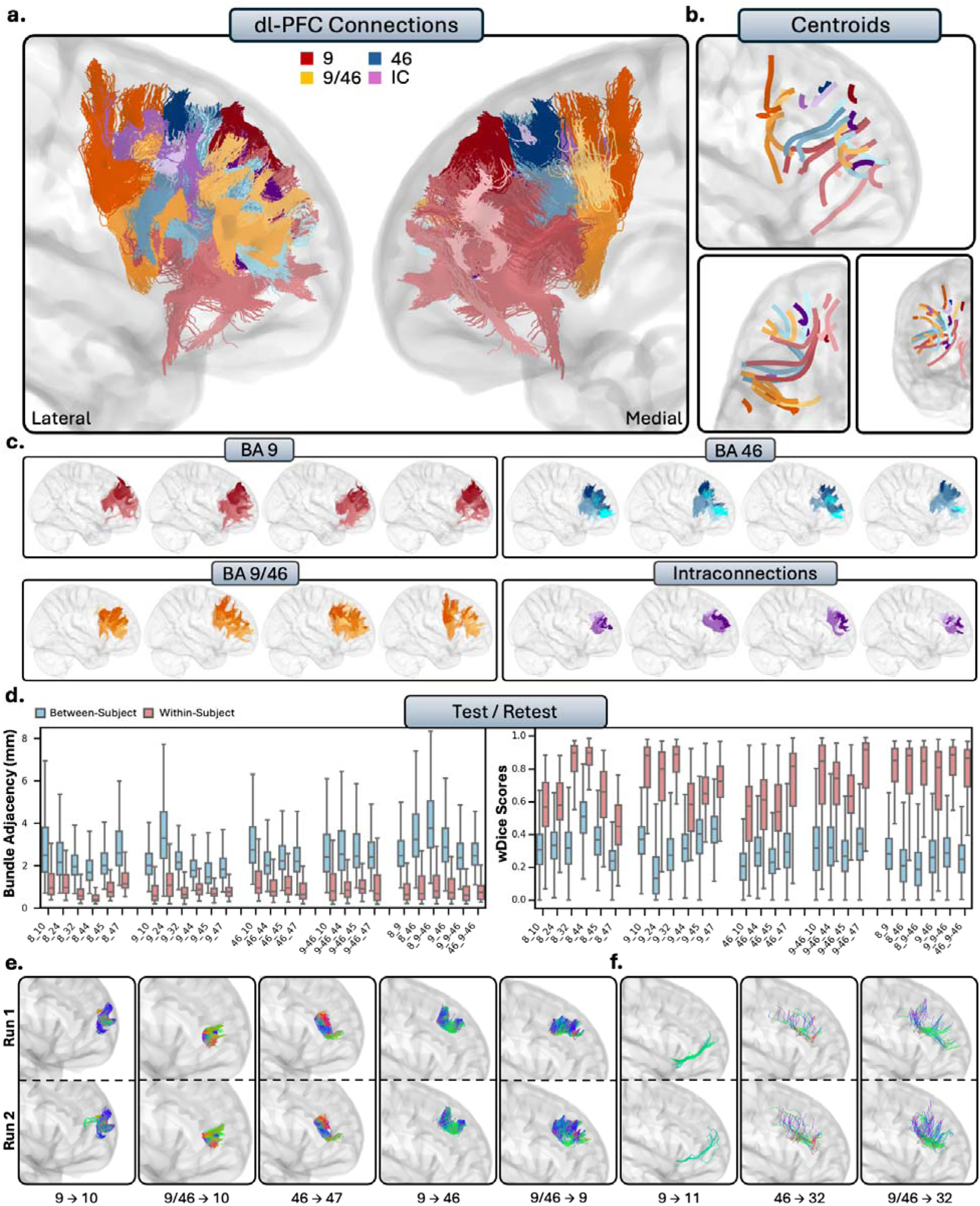
a.) The short connections of the dorsolateral prefrontal cortex (dl-PFC). Short connections from area 9 are colored red, area 46 are colored blue, 9/46 are colored orange, and interconnections (IC) are colored purple. Different shades indicate different connections. b.) Overall trajectories of the short fibers. Top: Sagittal view, left: axial view, right: coronal view. c.) Individual subject variability for each area in the dl-PFC. Each brain represents an individual subject’s trajectories from area 9, 46, 9/46 and the interconnections. For area 8, see Supplementary Figure 6. d.) Test-retest reliability of true positive tracts measured with bundle adjacency (left) and wDice (right). Blue bars indicate between-subject measurements and red bars indicate within-subject measurements. e.) Individual tract test-retest of representative RTP tracts and f.) examples of false positive tracts in the dl-PFC.

**Figure 4.**
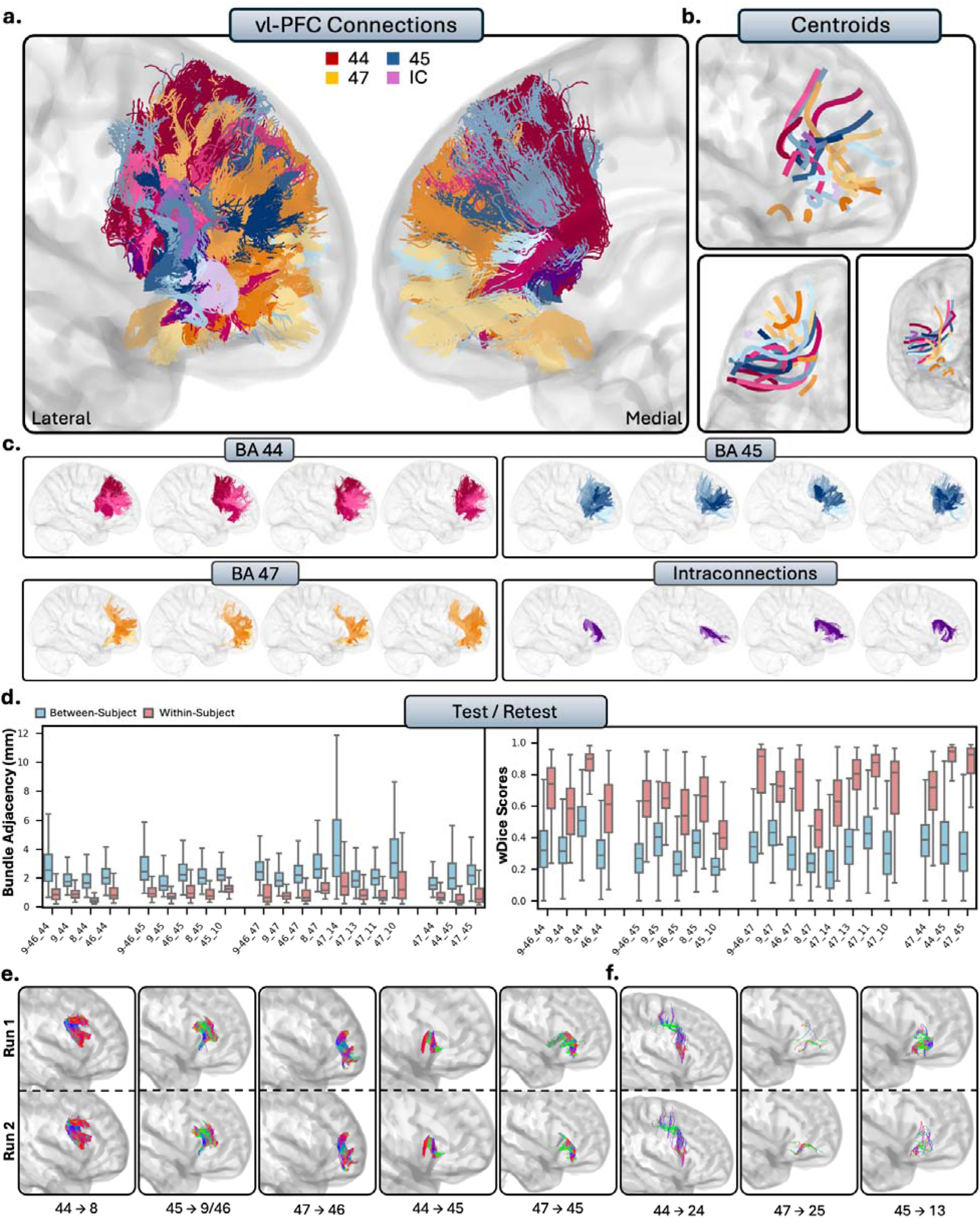
The short connections of the ventrolateral prefrontal cortex (vl-PFC). Short connections from area 44 are colored red, area 45 are colored blue, 47 are colored orange, and intraconnections (IC) are colored purple. Different shades indicate different connections. b.) Overall trajectories of the short fibers. Top: Sagittal view, left: axial view, right: coronal view. c.) Individual subject variability for each area in the vl-PFC. Each brain represents an individual subject’s trajectories from area 44, 45, 47 and the interconnections. d.) Test-retest reliability of true positive tracts measured with bundle adjacency (left) and wDice (right). Blue bars indicate between-subject measurements and red bars indicate within-subject measurements. e.) Individual tract test-retest of representative RTP tracts and f.) examples of false positive tracts in the vl-PFC.

**Figure 5.**
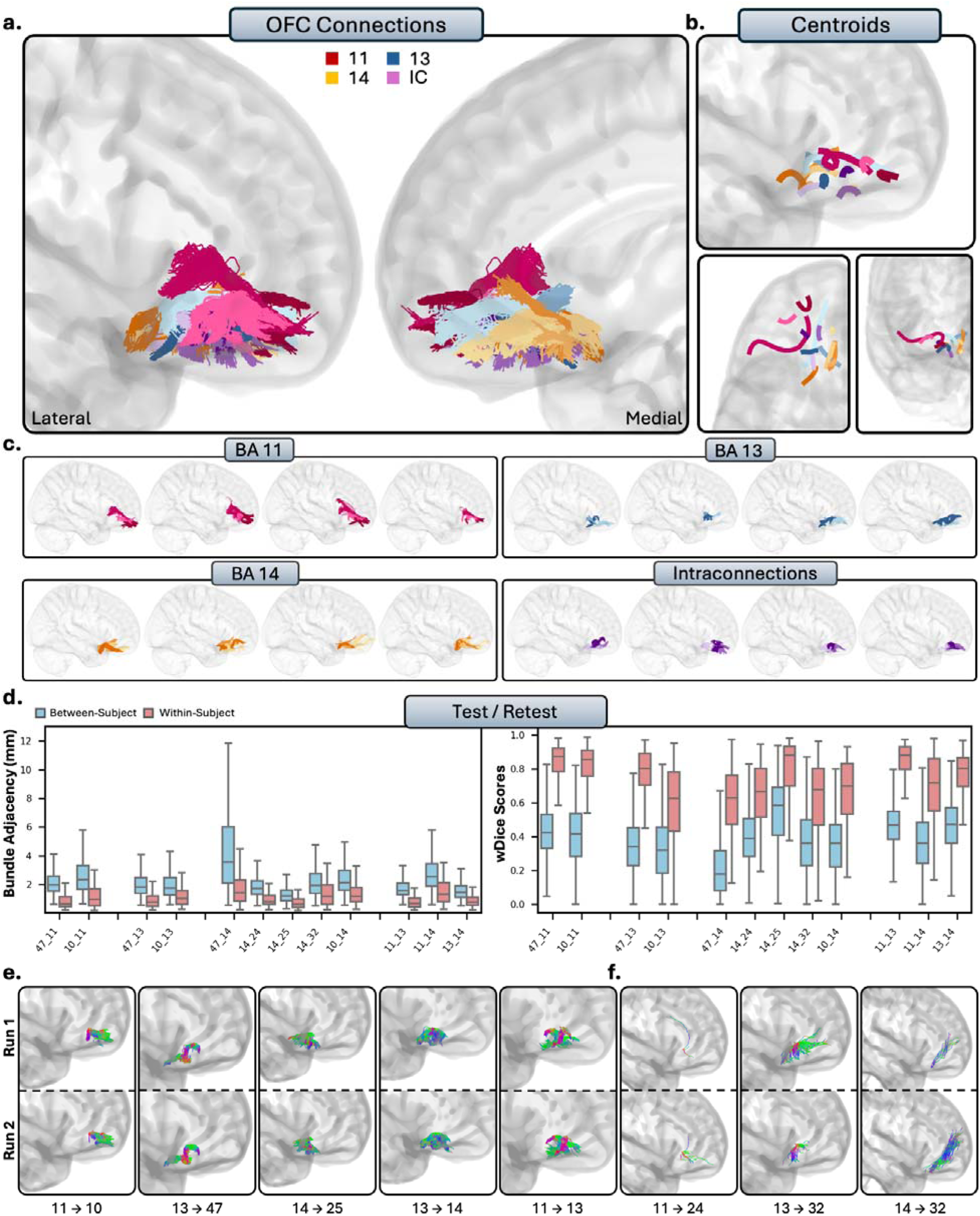
The short connections of the orbitofrontal cortex (OFC). Short connections from area 11 are colored red, area 11 are colored blue, 14 are colored orange, and interconnections are colored purple. Different shades indicate different connections. b.) Overall trajectories of the short fibers. Top: Sagittal view, left: axial view, right: coronal view. c.) Individual subject variability for each area in the OFC. Each brain represents an individual subject’s trajectories from area 11, 13, 14 and the interconnections. d.) Test-retest reliability of true positive tracts measured with bundle adjacency (left) and wDice (right). Blue bars indicate between-subject measurements and red bars indicate within-subject measurements. e.) Individual tract test-retest of representative RTP tracts and f.) examples of false positive tracts in the OFC.

**Figure 6.**
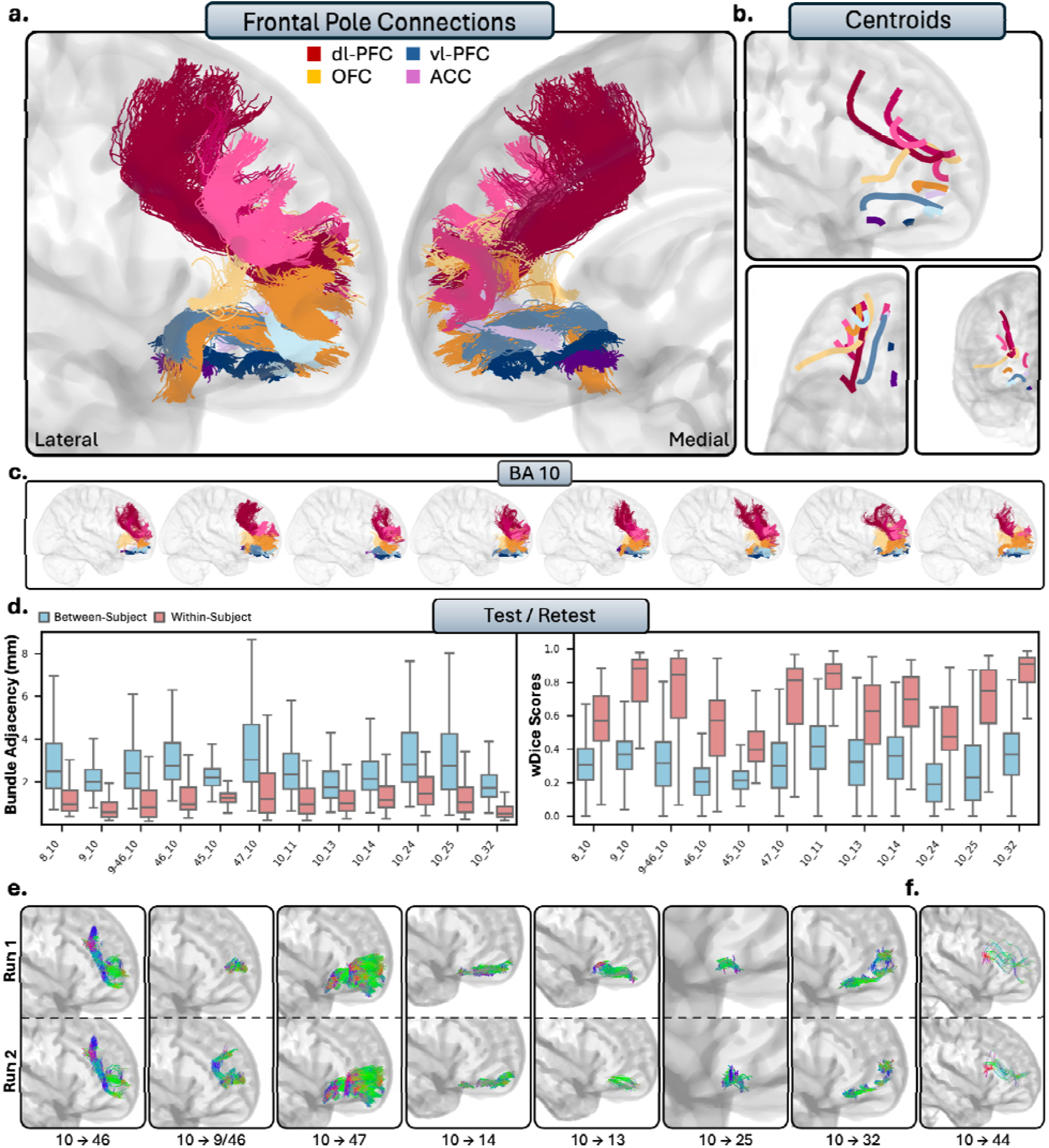
The short connections of the frontal pole. Connections to the dl-PFC are colored red, vl-PFC are blue, OFC are orange, and ACC are purple. Different shades indicate different connections. b.) Overall trajectories of the short fibers. Top: Sagittal view, left: axial view, right: coronal view. c.) Individual subject variability for each area in the frontal pole. Each brain represents an individual subject’s trajectories from each prefrontal region. d.) Test-retest reliability of true positive tracts measured with bundle adjacency (left) and wDice (right). Blue bars indicate between-subject measurements and red bars indicate within-subject measurements. e.) Individual tract test-retest of representative RTP tracts and f.) examples of false positive tracts between the frontal pole and area 44.

**Figure 7.**
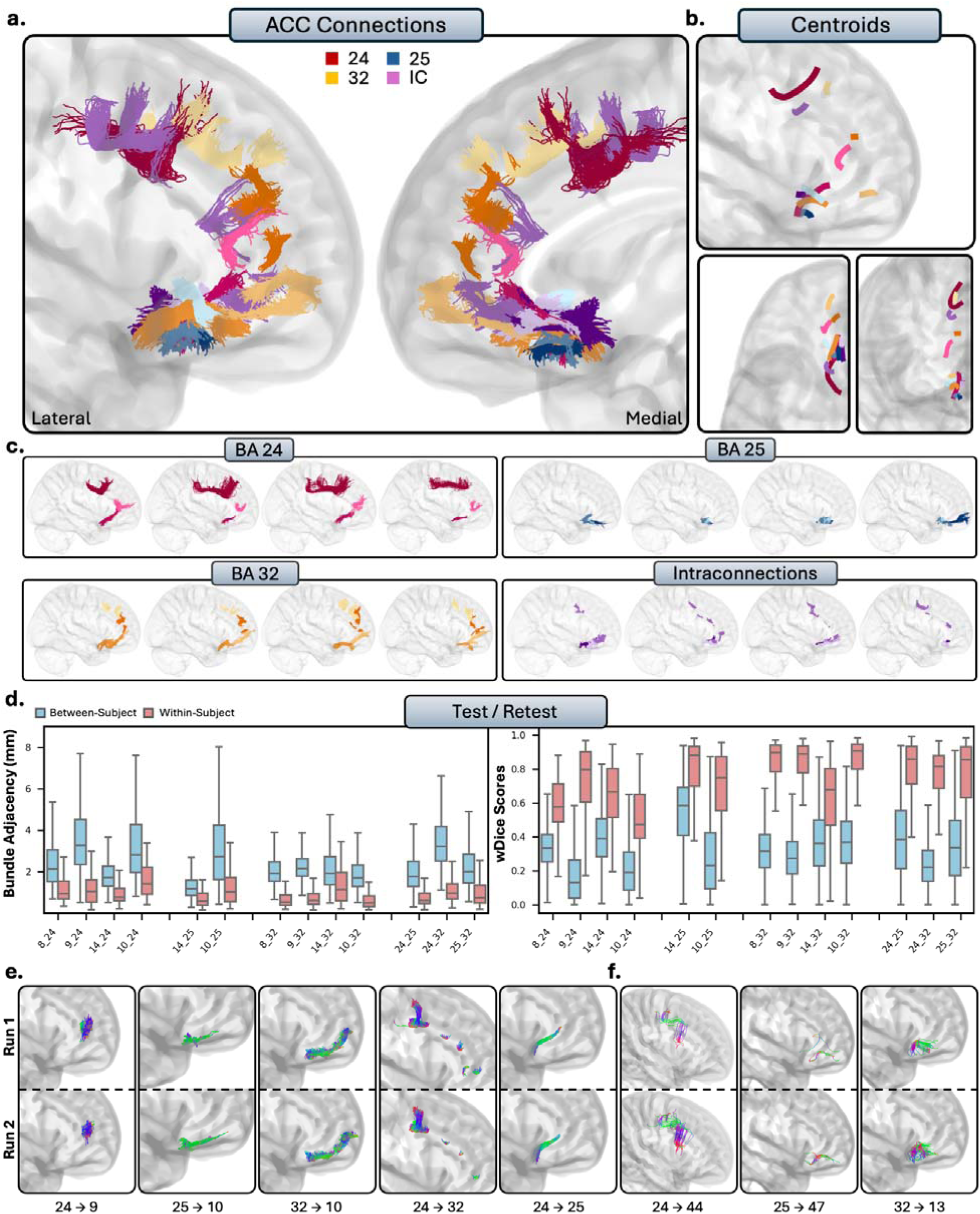
The short connections of the anterior cingulate cortex (ACC). Short connections from area 24 are colored red, area 25 are colored blue, 32 are colored orange, and interconnections are colored purple. b.) Overall trajectories of the short fibers. Top: Sagittal view, left: axial view, right: coronal view. c.) Individual subject variability for each area in the ACC. Each brain represents an individual subject’s trajectories from area 24, 25, 32 and the interconnections. d.) Test-retest reliability of true positive tracts measured with bundle adjacency (left) and wDice (right). Blue bars indicate between-subject measurements and red bars indicate within-subject measurements. e.) Individual tract test-retest of representative RTP tracts and f.) examples of false positive tracts in the ACC.

#### 3.2.1 Dorsolateral Prefrontal Cortex

The dl-PFC comprises Brodmann areas 8, 9, 46, and 9/46, which includes areas of the superior frontal gyrus (SFG) and the middle frontal gyrus (MFG). Qualitatively, the general patterns of connectivity were highly consistent across subjects (Fig. 3c-f). The short connections of the dl-PFC were characterized by relatively large, U-shaped bundles which were mostly localized to the superior and lateral portions of the hemisphere, with some exceptions where there was connectivity to the ACC. Note: Given the large volume of the connections involving area 8, we displayed them separately from the remaining dl-PFC pathways to enhance interpretability (Supplementary Figure 6).

##### Area 8

Area 8 corresponds to the dorsal and posterior portions of the SFG, reaching the MFG. It is one of the largest regions in this analysis, reaching over the medial wall of the sagittal plane and bordering the ACC. Area 8 showed connectivity with the vl-PFC: Area 8 and area 44 were connected with large bundles of U-shaped fibers which terminated throughout area 44 and several regions within area 8 (RTP). There were similar connections with areas 45 (TP) and 47 (TP), but were slightly less consistent across subjects compared to connections with 44. No connections were identified between area 8 and the OFC, consistent with histological reports for area 11 (TN) but inconsistent with histology for areas 13 (FN) and 14 (FN). There was connectivity between area 8 and the frontal pole, often consisting of two sub-bundles from the medial and lateral portions of area 8 (TP). The medial portions of area 8 showed connectivity to the ACC, but showed particularly close connectivity to area 32 (RTP). There was connectivity to area 24 across subjects (TP), and no connectivity to area 25, consistent with histological literature (TN). As areas 8, 32, and 24 are all quite large, each tract usually consists of multiple subdivisions throughout.

##### Area 9

Area 9 refers to the rostral superior frontal gyrus. It is bounded by the superior frontal sulcus, reaches over the sagittal plane, and borders the cingulate sulcus (Petrides et al., 2012). Overall, area 9 showed connectivity to all regions of the vl-PFC, with multiple bundles of large, U-shaped fibers stemming from both the lateral and medial portions of area 9 to areas 45 (RTP) and 47 (RTP). Connections to area 44 were more sparse and sometimes contained spurious streamlines, but were consistent and anatomically plausible across the majority of subjects (TP). There were no connections between area 9 and the OFC: there were no connections with and 14 despite histological precedence (FN), and only false positive connections with the areas 11 (FP) and 13 (FP). Area 9 showed close connectivity with the frontal pole, often with multiple subdivisions (RTP). This is particularly true with rostral area 9, often displaying several U-shaped bundles which follow the gyral folds of the rostral SFG.

Medial area 9 was connected with areas 24 (RTP) and 32 (RTP) of the ACC. The connections between area 9 and 24 were small, dense bundles of streamlines travelling vertically between 9 and anterior 24 and on occasion streamlines would reach more posterior portions of 24. There were similarly small, dense connections between area 9 and 32, with several subdivisions along the rostral/caudal plane. There were no connections between areas 9 and 25, consistent with histological literature (TN).

##### Area 46

Area 46 refers to the central middle frontal gyrus, and is both anteriorly and posteriorly bounded by area 9/46. Area 46 showed connectivity with the vl-PFC: connections between areas 44 and 46 were dense, U-shaped bundles (RTP). Connections between areas 8 and 46 were dense as well (RTP), oftentimes displaying two subdivisions of these U-shaped bundles, with one travelling more anterior to posterior, and one travelling more dorsal to ventral. The connections between areas 46 and 47 were typically dense U-shape between the two regions (RTP). There were dense connections to area 45, but sometimes contained spurious tracts in the posterior portion of the bundles (TP). No tracts were modeled between area 46 and the OFC, consistent with histological literature (11, 13, 14 TN). Area 46 showed connectivity to the frontal pole (TP). There were no connections between areas 46 and 24, despite histological precedence (FN). There were also connections with area 25, which is consistent with histological literature (TN). There were spurious, anatomically implausible connections between areas 46 and 32 across the majority of subjects, sometimes including portions of the corpus callosum (FP).

##### Area 9/46

Area 9/46 is also located in the middle frontal gyrus. Proposed in 1999 by Petrides and Pandya, and is typically subdivided in the literature into two portions, dorsal 9/46 and ventral 9/46 (Petrides & Pandya, 1999). Area 9/46 showed dense connections with the vl-PFC, with connections to areas 44 (RTP), 45 (RTP), and 47 (RTP). The connections to areas 45 were relatively large, often stemming from both the dorsal and ventral portions of 9/46, whereas area 44 connections mostly stemmed from the ventral portion of 9/46. Connections with area 47 were smaller in volume, but were very dense, and mostly with the ventral portion of 9/46. There were no modeled connections between area 9/46 and the OFC despite histological precedence, with no modeled connections between areas 9/46, 13 (FN), and 14 (FN), and only few spurious connections inconsistently found across subjects between areas 9/46 and 11 (FN). There were dense, U-shaped connections between areas 9/46 and the frontal pole, typically stemming from the anterior portions of ventral 9/46 (RTP). No connections were detected between area 46 and ACC areas 24 (FN) and 32 (FN), despite histological evidence for these pathways. In contrast, the absence of connections between areas 9/46 and area 25 (TN) was consistent with histological literature.

##### Intraconnections

The dl-PFC was highly intraconnected, as areas 8, 9, 46, and 9/46 all displayed dense reciprocal connections to each other (RTP). The intraconnections of the dl-PFC were characterized by multiple, dense, U-shaped subdivisions within the tracts, and were highly variable between individuals. The connections between areas 8 with 9, 8 with 9/46, and 9 with 9/46 all consistently showed two or more subdivisions within the bundles, though these subdivisions were present in most intraconnections of the dl-PFC.

##### Summary

The dl-PFC overall had strong connectivity with the vl-PFC and the frontal pole, as all areas showed connections to both regions, with exception of area 8. The dl-PFC showed moderate connection with the ACC, with the medial portions of areas 8 and 9 showing connections with BA 32 in particular. Interestingly, there were no modeled connections between the dl-PFC and the OFC.

#### 3.2.2 Ventrolateral Prefrontal Cortex

The vl-PFC comprises areas 44, 45, and 47, which correspond to the pars opercularis, pars triangularis, and pars orbitalis of the inferior frontal gyrus, respectively. While Area 47 is included in the vl-PFC for this analysis, it is sometimes considered part of the OFC due to its proximity to the orbital gyri. Qualitatively, the vl-PFC connections were characterized by large, lateral, U-shaped bundles (Fig. 4c-f). Unlike the dl-PFC, the tracts from the vl-PFC in general were much more lateral among subjects, as few connections reach the ACC. The connections of vl-PFC were notably anteroventral and displayed much closer connections to the OFC than other regions of the PFC.

##### Area 44

Area 44 refers to the opercular portion of the IFG, and is often referred to as Broca’s area. Overall, area 44 showed limited connectivity to the PFC. Area 44 was connected with the dl-PFC, with dense connections to areas 46 (RTP) and the posterior ventral portions of 9/46 (RTP). The connections between area 44 and area often exhibited two large, dense subbundles stemming from both the medial and lateral portions of area 8, terminating in the entire surface of area 44 (RTP). Connections with area 9 were volumetrically large, dense, U-shaped bundles of streamlines. However, they were not as consistent across the population compared to its connections with the other regions of the dl-PFC (but still remained TP). Consistent with histological reports, area 44 and the OFC showed no connectivity, as there were sparse inconsistent streamlines with area 11 (TN), and no streamlines modeled at all with 13 (TN) or 14 (TN). Similarly, there were few but inconsistent, spurious tracts between area 44 and the frontal pole (TN), consistent with histology. Consistent with histology, no tracts were present between areas 25 (TN) of the ACC. There were consistent, spurious tracts with areas 24 (FP) and 32 (FP).

##### Area 45

Area 45 displayed connectivity with all areas of the dl-PFC. Areas 9 (RTP) and ventral 9/46 (RTP) had large U-shape connections that travelled ventrally and terminated in the entire surface of area 45, consistent across the entire population. Area 45 had variable connectivity with areas 8 (TP) and 46 (TP): oftentimes connections were dense and anatomically plausible, but at times contained false positives due to large numbers of spurious streamlines.

There was less connectivity within the OFC. There were often tracts that seemed anatomically plausible reaching from the anterior portions of area 45 to the superior portions of areas 11, but the majority of subjects displayed clear false positive bundles (FP). However, no tracts were found with area 13 despite histological evidence (FN), while the lack of connections with area 14 was consistent with histology (TN). There were connections from area 45 to the frontal pole: they often were large in volume and dense (TP), but had the tendency to include spurious streamlines. For the ACC, no tracts were present between areas 24 (FN), and 25 (TN), and anatomically implausible, consistent bundles between areas 45 and 32 (FP).

##### Area 47

Area 47, often referred to as area 47/12 in the macaque literature, refers to the pars orbitalis of the IFG and lateral portions of the orbital gyri. Area 47 was connected with the dl-PFC, with dense connections between areas 9 (RTP), 9/46 (RTP), and 46 (RTP), with similar but slightly less consistent connections with area 8 across the population (TP). Connections with area 9 were volumetrically large and often contained two subbundles, stemming from either dorsomedial area 9 and reaching to dorsolateral 47, or stemming from ventrolateral 9 and reaching to more orbital portions of 47. Area 46 and 47 displayed particularly close, dense, U-shaped connections. Connections with ventral area 9/46 were small, dense, and highly individualized with multiple subbundles. Unsurprisingly, area 47 was connected to all regions of the OFC, with consistent connectivity with areas 11 (RTP), 13 (RTP), and 14 (RTP). These bundles typically had multiple small, dense, U-shape connections which traveled from the orbital portions of area 47. In particular, area 11 displayed short-range dense connections with orbital 47, which sometimes reached the lateral portions of area 47 (RTP). The connections between area 47 and the frontal pole were dense and widely individually variable (RTP). There were often two separate bundles, one stemming from the lateral portions of 47 to the superior portions of area 10, and one stemming from the orbital portions of 47 to the anteroventral portions of area 10. There were no connections between area 47 and the ACC, as no tracts were present between area 24 (FN), which differs from histology. There were also no bundles with 25 (TN), which was consistent with histology. The majority of tracts modeled between area 47 and 32 were spurious and were not anatomically plausible (FP). There were subjects (ex., subjects 656253, 702133, 908860) which displayed plausible connections between areas 32 and the most orbitomedial portions of 47. However, these bundles were reconstructed in less than 50% of subjects.

##### Intraconnections

The vl-PFC was highly intraconnected, as areas 44, 45, and 47 all displayed reciprocal connections to each other (RTP). Areas 44 and 45 in particular were closely connected, forming a dense U-shape as the streamlines followed the gyral folds between the pars opercularis and pars triangularis. There were often two subbundles between 44 and 45, one which closely followed the gyral folds, as well as one which formed a U-shape and travelled more medially into the white matter.

##### Summary

The vl-PFC is highly interconnected with the rest of the PFC, but not to the same extent as the dl-PFC. The vl-PFC shares high levels of connectivity with the dl-PFC and the frontal pole, with all areas of the vl-PFC displaying some connectivity to each region. Area 47 had the most connectivity throughout the PFC, particularly with the OFC, displaying close connections with all areas within the OFC. However, we found very limited connectivity between vl-PFC and the ACC.

#### 3.2.3 Orbitofrontal Cortex

The OFC comprises areas 11, 13, and 14, and is located on the ventral surface of the PFC. The general organization of the OFC was conserved between individuals: tracts were notably more compact, densely packed, and predominantly localized to regions bounded by the OFC. Tracts were much smaller in volume when compared to other regions, and typically formed small, condensed U-shapes as they traversed the orbital gyri.

##### Area 11

Area 11 showed no connectivity to the dl-PFC. Consistent with histological literature, there were no connections present between area 46 (TN) or area 8 (TN). Conversely, there were anatomically implausible streamlines with areas 9 (FP) and 9/46 (FP) consistent across the population. Area 11 showed inconsistent connectivity with the vl-PFC, with no tracts passing visual inspection between 11 and 44 (TN), consistent with histological literature. There were mostly anatomically implausible bundles reconstructed between areas 11 and 45 (FP), despite some subjects displaying anatomically plausible bundles. The connections between areas 11 and 47 were among the densest and most consistent connections between area 11 and the rest of the PFC, barring the interconnections between the OFC and frontal pole (RTP). These connections formed small, dense U-shaped connections from area 11 to the medial portions of area 45, often stretching to the lateral portions. The connections between area 11 and the frontal pole consisted of small, U-shaped tracts extending anteromedially from area 11 to the most anterior regions of the frontal pole (RTP). Area 11 showed no connectivity to the ACC. Areas 24 showed no tracts, despite histological precedence (FN). Consistent with histology, there were now tracts with area 25 (TN). There were only spurious tracts between 11 and 32 (FP).

##### Areas 13 and 14

Within the HCP-MMP1 atlas, areas 13 and 14 share many regions (see Table 1). As a result, areas 13 and 14 show very similar patterns of connectivity, with few key differences. Areas 13 and 14 showed no connectivity to the entire dl-PFC, despite histological evidence for areas 8 (13 = FN, 14 = FN), 9 (13 = FN, 14 = FP), and 9/46 (13 = FN, 14 = FN). However, this was consistent with histology for area 46 (13 = TN, 14 = TN). For connections with the vl-PFC, areas 13 and 14 modeled no tracts between areas 44 (TN, TN), consistent with histological literature. For area 45, there were no tracts, differing with histology for area 13 (FN), but consistent for area 14 (TN). However, similar to area 11, areas 13 (RTP) and 14 (RTP) showed dense, short-range connections to area 47. Connections between areas 13 and 47 were typically very densely packed U-fibers, traversing from the lateral portions of 13 to the medial portions of 47. In contrast, connections between areas 14 and 47 stemmed from more posteromedial portions of the OFC to area 47, and often reach the most anterior portions of the orbital surface of area 47. Connections from areas 13 (RTP) and 14 (RTP) to the frontal pole were similar, each consisting of dense tracts that travelled posterior-to-anterior from the OFC to the frontal pole, with connections from area 14 being more medial. The ACC is where areas 13 and 14 differed the most. No connections were present between area 13 and areas 24 (FN) and 25 (FN) despite histological evidence, and only spurious bundles with area 32 (FP). Area 14 displayed mostly consistent connectivity to area 24 (TP), and dense short-range connectivity to areas 25 (RTP) and 32 (RTP).

##### Intraconnections

The OFC was highly intraconnected, as areas 11, 13, and 14 all displayed dense reciprocal connections to each other (RTP). Area 11 showed clear, dense patterns of connectivity to both areas 13 and 14. Although connections between areas 13 and 14 were robust, they were more difficult to disentangle due to overlapping space. Multiple subbundles were often present, typically comprising a U-shaped posterior tract and a linear anterior-to-posterior tract stemming from the anterior portion of area 14.

##### Summary

Compared to the rest of the PFC, the connections of the OFC were more localized, and displayed less widespread intercortical connectivity. Connectivity to the dl-PFC was absent, and connections to the vl-PFC were relatively sparse, except for robust connections throughout the OFC with area 47. The OFC also showed notable connections with the frontal pole. Area 14, but not areas 11 or 13, displayed notable connectivity with the ACC. Within the OFC, areas 11, 13, and 14 were highly intraconnected.

#### 3.2.4 Frontal Pole

Unlike the other regions of the PFC, the frontal pole consists of only one Brodmann area, BA 10. The frontal pole is the most anterior portion of the PFC, dorsal to the OFC and anterior to the dl-PFC and vl-PFC. The tracts from the frontal pole were consistently modeled in each individual, but as it is a large area, there was a significant amount of individual variability.

Nonetheless, the tracts reaching the frontal pole qualitatively displayed a consistent organization: large, elongated U-shaped fibers originating from the posterior PFC, and shorter, densely packed fibers arising from the anterior PFC. Notably, the frontal pole was interconnected with the rest of the PFC: the frontal pole exhibited 12 true positives (9 RTP) and 1 true negative, making the frontal pole the region with the most histological accuracy at a perfect 100%.

##### Connections with dl-PFC

The frontal pole was connected to the dl-PFC, with consistent connections to areas 8, 9, and 9/46, and TP connections to area 46. Frontal pole connections to area 8 were often subdivided into two subbundles, stemming from either the medial or lateral portions of area 8. Similarly, there were often many subbundles within the connections of area rostral 9 and the frontal pole. Connections with area 9/46 were typically dense, U-shaped bundles, and were mostly connected to the ventral portion of 9/46. Area 46 connections were consistently created between individuals, but sometimes included false positive tracts. Nonetheless, tracts from area 46 to the frontal pole were generally large, U-shaped connections, travelling medially and rostrally to the rostral portions of the frontal pole.

##### Connections with vl-PFC

The frontal pole showed connectivity with the vl-PFC, with exception of area 44, where sparse anatomically implausible streamlines were sometimes present (TN), consistent with literature. Area 45 and the frontal pole often exhibited large, wide U-shaped fibers which typically reached the lateral portions of the frontal pole from area 45 (TP). The connections between the frontal pole and area 47 were very dense, robust connections, which were consistently replicated between subjects. The connections from area 47 typically reached the anteroventral portions of the frontal pole, and were relatively large in volume (RTP).

##### Connections with OFC

The frontal pole exhibited robust, consistent connectivity to the entire OFC. Area 11 and the frontal pole were typically connected by small, consistent, and dense U-shaped bundles, which reached the anterolateral tip of the frontal pole (RTP). Connections with area 13 were dense, short projections which reached the lateral frontal pole (RTP), and connections from area 14 were small, U-shaped connections that typically reach that medial surface of the frontal pole (RTP).

##### Connections with ACC

The frontal pole and the ACC displayed differing connectivity between each area. There were consistent, dense connections between the frontal pole and rostral area 24 (RTP). These were anatomically viable, however, at times these tracts would mix with invalid streamlines from corpus callosum. Area 25 and the frontal pole consistently demonstrated dense, volumetrically thin tracts that stemmed from area 25 and reached to the posterior portions of the frontal pole (RTP). Lastly, there was robust, consistent connectivity between the frontal pole and area 32 (RTP). Typically, these connections were multiple small, dense subbundles that connected the anteroventral area 32 to the ventral and middle frontal pole.

##### Summary

The frontal pole was highly interconnected with the rest of the PFC. It showed robust connections with areas 8, 9, and 9/46 of the dl-PFC, with additional but less consistent connections to area 46. Within the vl-PFC, the frontal pole was strongly connected to areas 45 and 47, but not to area 44. It also displayed consistent connections with OFC areas 11, 13, and 14. Lastly, the frontal pole showed variable connectivity with the ACC, with connections to 25 and 32, and variable true positives with 24.

#### 3.2.5 Anterior Cingulate Cortex

The anterior cingulate cortex comprises areas 24, 25, and 32, and is located on the medial surface of the brain on the cingulate gyrus. While general patterns of connectivity were conserved between subjects, the ACC displayed the most clear individual variability, as tracts from the ACC were typically groups of small, U-shaped bundles which followed the cingulate gyrus.

##### Area 24

Area 24 of the ACC is a large region of grey matter that bounds the genu of the corpus callosum, posterior to area 32, and displayed limited connectivity to the rest of the PFC. Area 24 showed connectivity with the dl-PFC, as there was connectivity to medial area 8 (TP), and dense, consistent groups of bundles with anteromedial, and occasionally reaching posteromedial area 9 (RTP). Despite histological reports, there were no valid connections with area 46 (FN), and only spurious tracts with area 9/46 (FP). There were no valid connections between area 24 and the vl-PFC, with only spurious, anatomically implausible bundles in area 44 (FP), and no tracts present in areas 45 (FN) and 47 (FN), differing from histological studies. Within the OFC, areas 11 (FN) and 13 (FN) showed no connectivity to area 24. Area 14 displayed small, dense anterior-to-posterior connections with area 24, with some consistency (TP). The frontal pole consistently displayed tracts to area 24 between subjects (TP), but occasionally would mix invalid tracts which appeared to stem from the corpus callosum.

##### Area 25

Area 25 is a very small region ventral to the genu of the corpus callosum. Area 25 was the least interconnected region of the PFC, as there were no valid connections present with either the dl-PFC (TN) or the vl-PFC (TN). Area 25 displayed variable connectivity to the OFC, depending on the region. No tracts were present with area 11 (TN), consistent with histological reports. There were small and dense bundles between areas 25 and 13, though in fewer than half of the subjects (FN). Additionally, area 14 exhibited robust, dense connectivity with area 25 (RTP). The frontal pole also showed consistent connectivity to area 25, with dense, volumetrically thin tracts extending from area 25 to the posterior portions of the frontal pole (RTP).

##### Area 32

Area 32 is a large segmentation of the grey matter in the anterior portion of the cingulate cortex, anterodorsal to area 24. Area 32 exhibited broader and more consistent connectivity with the PFC compared to areas 24 and 25. In the dl-PFC, the medial portions of areas 8 (RTP) and 9 (RTP) displayed dense bundles connecting to area 32, with area 9 in particular exhibiting dense bundles of varying volume along the anterior-posterior plane. Only anatomically implausible streamlines were reconstructed between areas 46 (FP) and 9/46 (FP), with connections to area 46 often including mixing with streamlines stemming from the corpus callosum. In the vl-PFC, only anatomically implausible tracts were reconstructed between areas 44 (FP), 45 (FP), and 47 (FP). However, a few subjects exhibited viable connections between orbitomedial 47 and area 32, though these were infrequent. Within the OFC, areas 11 (FP) and 13 (FP) showed only false positive bundles with area 32. Area 14 demonstrated robust, dense connectivity with the posteroventral portions of area 32 (RTP). Lastly, the frontal pole displayed robust, consistent connections to area 32, with multiple small, dense subbundles linking anteroventral area 32 to the ventral and middle frontal pole (RTP).

##### Intraconnections

Like other regions of the PFC, the ACC was highly intraconnected, as areas 24, 25, and 32 displayed dense reciprocal connections (RTP). Connections from area 24 to 25 were typically thin, densely packed bundles along the rostral-caudal which connected the two areas at the anteroventral portions of area 24. Connections between areas 24 and 32 consisted of many smaller dense subbundles along the bounding areas of the two regions as they run in tangent with each other. Lastly, connections between 25 and 32 were small, dense connections which reached the most posteroventral portions of 32. However, there were a few subjects who failed visual inspection, as some tracts erroneously mixed with the anterior commissure.

##### Summary

The ACC exhibited fewer connections in comparison to the rest of the PFC. Connectivity to the dl-PFC was limited, only displaying connections with areas 8 and 9. Connections to the vl-PFC were functionally absent, with no areas of the ACC showing viable streamlines, besides occasional connectivity between 32 and 47. In the OFC, area 14 had consistent connectivity with the ACC. Lastly, the frontal pole showed variable connectivity with the ACC, with valid connections to areas 25 and 32, and variable connections to area 24.

### 3.3 Test-Retest Reliability

We measured test-retest reliability in a subsection of subjects who had two sets of diffusion and structural scans (Figs. 3-7d). Within- and between-subject metrics for each PFC subregion are fully reported in Table 2. Within-subject reliability was high, as between scans spatial disagreement was exceptionally low (mean BA range = 1.01mm - 1.35mm), and volumetric overlap was moderate to high (mean wDice range = 0.67 - 0.73). Conversely, between-subject spatial disagreement was much higher (mean BA range = 2.28mm - 2.71mm) and volumetric overlap was much lower (mean wDice range = 0.30 - 0.39). This pattern was consistent for each partition of the PFC, and suggests high scan-rescan reliability within-subjects, but substantial individual differences across the population.

**Table 2:** Overview of Test-Retest Reliability.

| **PFC Partition** | **Spatial Disagreement (BA)** |  | **Volumetric Overlap (wDice)** |  |
| --- | --- | --- | --- | --- |
|  | Within-Subject | Between-Subject | Within-Subject | Between-Subject |
| dl-PFC | $1.01 \pm 0.83\text{mm}$ | $2.66 \pm 1.41\text{mm}$ | $0.70 \pm 0.22$ | $0.30 \pm 0.15$ |
| vl-PFC | $1.05 \pm 0.95\text{mm}$ | $2.49 \pm 1.49\text{mm}$ | $0.69 \pm 0.22$ | $0.33 \pm 0.16$ |
| OFC | $1.23 \pm 1.12\text{mm}$ | $2.28 \pm 1.53\text{mm}$ | $0.72 \pm 0.22$ | $0.39 \pm 0.18$ |
| Frontal Pole | $1.35 \pm 1.37\text{mm}$ | $2.71 \pm 1.69\text{mm}$ | $0.67 \pm 0.24$ | $0.32 \pm 0.17$ |
| ACC | $1.12 \pm 1.04\text{mm}$ | $2.44 \pm 1.41\text{mm}$ | $0.73 \pm 0.22$ | $0.33 \pm 0.19$ |

Measurements represent mean values calculated across pathways, ± = standard deviation; wDice = weighted dice; BA = bundle adjacency; dl-PFC = dorsolateral prefrontal cortex; vl-PFC = ventrolateral prefrontal cortex; OFC = orbitofrontal cortex; ACC = anterior cingulate cortex

## 4. Discussion

Here, we present the first systematic *in-vivo* visualization of the short-range connections of the prefrontal cortex in the human brain. Guided by decades of histological work, our analysis revealed three takeaways: (1) short cortico-cortical pathways can be reliably reconstructed, showing strong histological correspondence and robustness to false positives; (2) these connections are highly stable within individuals but exhibit substantial variability across the population, revealing a unique connective fingerprint; and (3) reconstruction success varies by region, with lateral and frontal pole areas showing higher fidelity than medial and orbital regions. In the following sections, we discuss the implications of these findings for the study of the human connectome.

### Visualization of Short Association Fibers

The SAFs of the human brain, particularly in complex association areas, are underexplored, as their small size, variability, and partial volume effects make them challenging for tractography (Schilling et al., 2025). Although SAFs have received increasing attention in recent years, they are a relatively young area of research, and no standardized framework exists for their investigation. Prior studies have examined SAFs throughout the brain using a range of methodological approaches (Guevara et al., 2020). One commonly used strategy most similar to the present work employs atlas-derived ROIs to perform targeted tractography between selected regions, either at the whole-brain level (Nazeri et al., 2015) or within specific systems such as the supplementary motor area (Bozkurt et al., 2016), visual processing pathways (Movahedian et al., 2025), and short-fiber networks spanning frontal, temporal, and parietal cortices (Shukla et al., 2011). Another common approach uses hand-placed anatomically informed ROIs to reconstruct specific sets of short-range fibers. A seminal example is the work by Catani and colleagues (2012), who systematically reconstructed frontal SAFs using spherical ROIs placed along major gyri and validated these pathways with post-mortem dissection. Variants of this strategy have since been applied beyond the frontal lobe, including temporal, occipital, and parietal regions (Wu et al., 2016; Burks et al., 2017; Catani et al., 2017). More recently, fully data-driven clustering methods have been introduced, grouping superficial streamlines based on geometric similarity and spatial proximity to identify bundles either within specific regions or across the entire tractogram (Guevara et al., 2017; Yeh et al., 2018; Zhang et al., 2018; Zhang et al., 2019b; Guevara et al., 2020; Román et al., 2022).

Collectively, these studies suggest that SAFs can be reliably reconstructed with high anatomical fidelity when tractography is grounded in biological constraints (Schilling et al., 2020), particularly methods using histology-informed ROI placement (De Schotten et al., 2011; Rolls et al., 2023; Amandola et al., 2025), or which have been validated using Klinger dissection (Catani et al., 2012; Burks et al., 2017). Building on this foundation, we used histological findings to visualize SAFs across the entire PFC. We found that these short connections are measurable with tractography, as we were able to model the SAF’s of the PFC at an accuracy of 74%, driven by a high sensitivity of 81% with respect to past histological literature (Fig. 8). Though these are small, complicated bundles, our methodology seems to perform comparably with well-established deep white matter association fibers in NHP and humans (Knösche et al., 2015; Maier-Hein et al., 2017). Performance was highest in the lateral portions of the PFC, as the dl-PFC, vl-PFC, and particularly the frontal pole performed well compared to the ACC and medial portions of the OFC, likely a result of low signal-to-noise ratio in the OFC (Koch et al., 2002) and medial PFC (Koch et al., 2002; Clark et al., 2021), and interference from crossing fibers such as the cingulum and CC (Haber et al., 2022).

**Figure 8.**
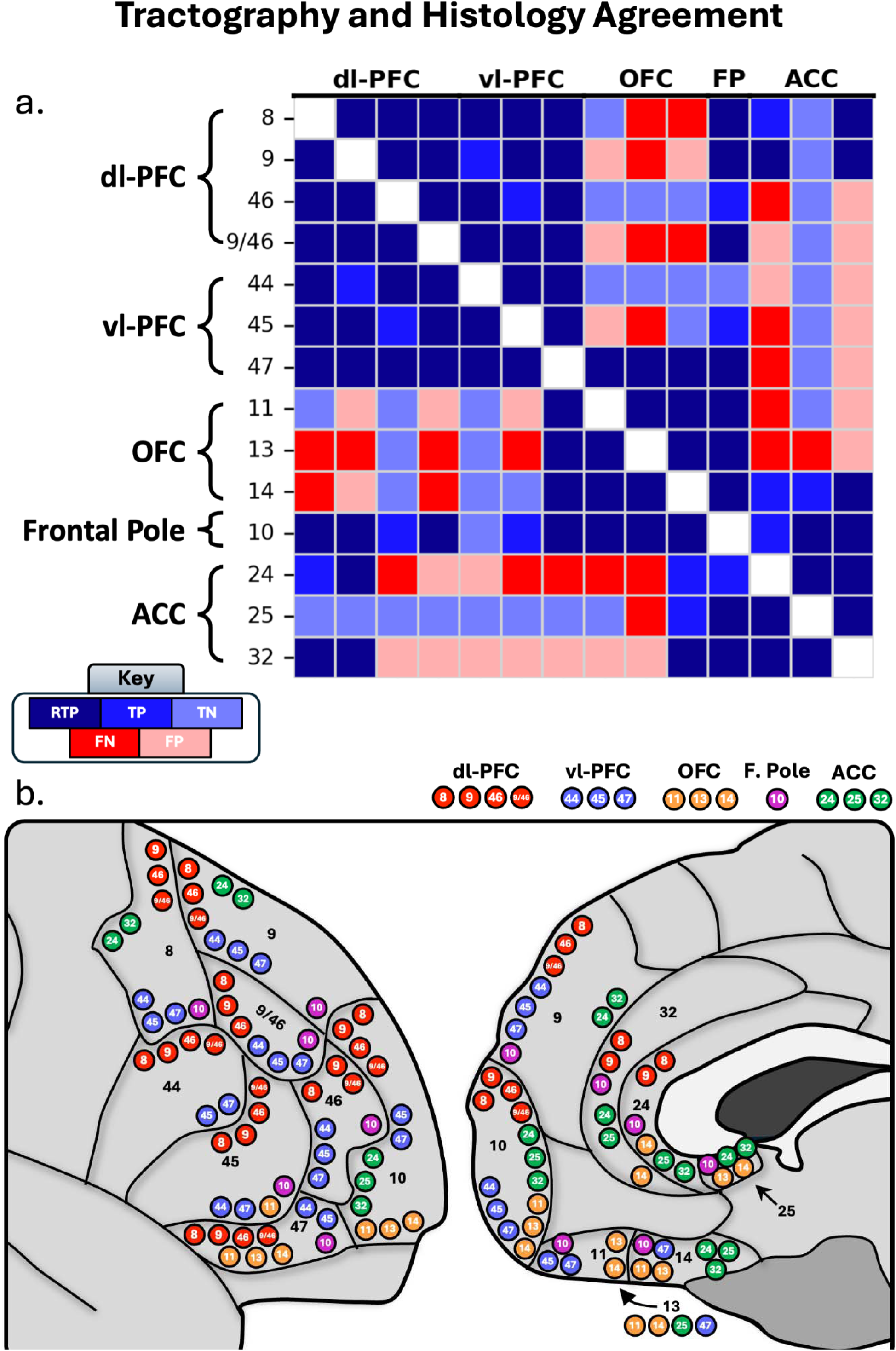
a.) Correspondence between tractography results and histology literature. Dark blue corresponds to robust true positives (RTP), blue corresponds to true positives (TP), light blue corresponds to true negatives (TN). Dark red corresponds to false negatives, and light red corresponds to false positives. b.) Schematic depicting the True Positive connections where tractography and histology converge. Red = dl-PFC, blue = vl-PFC, orange = OFC, purple = frontal pole, green = ACC.

Our results also demonstrated notable precision. While tractography can often reproduce known positive connections, this typically occurs at the cost of specificity (Thomas et al., 2014; Knösche et al., 2015; Maier-Hein et al., 2017; Schilling et al., 2019). Indeed, prior work suggests that connectomes may contain up to four times as many invalid as valid bundles, yielding an average precision of only 23% (Maier-Hein et al., 2017). In contrast, our approach achieved a precision of 79%, modeling 49 true positive and only 13 false positive tracts. Moreover, our method was more sensitive to true negatives than to false positives, with a specificity of 57% (for overview of false negative and false positive tracts, see Supplementary Figure 7). Recently, Girard and colleagues (2020) conducted a similar tractography study in an ex-vivo rhesus macaque brain where they evaluated the concurrence of connections created by tractography with frontal and parietal connections found in histological literature. While this study was not explicitly testing the SAFs of the PFC, all connections between the 59 cortical regions were evaluated with tractography and found that the short-range fibers displayed particularly high concurrence (Girard et al., 2020). This is an especially elucidating study, as it parcellates the prefrontal regions beyond full Brodmann Areas (i/e. 8Ad, 8Av, 8B vs. Area 8), and shows tractography’s ability to reconstruct these even smaller, more subdivided bundles. While our results are not intended to be directly or perfectly concurrent with those reported by Girard and colleagues (2020), this work nonetheless increases confidence in tractography’s ability to reconstruct short-range frontal connections consistent with known primate anatomy. Together, in combination with our results, this indicates that anchoring tractography in the biological framework established by histological research enables reconstruction of a more anatomically plausible and specific map of the short-range human connectome.

### Replication of Histological Findings

A central goal of this study was to determine if our histology-informed approach could replicate the organizational principles of PFC connectivity established by decades of primate tract-tracing. Overall, our tractography findings demonstrate a strong correspondence, capturing not only broad organization rules but also detailed, region-specific patterns only seen in invasive studies.

One of the most replicable findings in the histological literature is that adjacent areas share strong connectivity with each other (Yeterian et al., 2012, Haber et al., 2022). This property of adjacent connectivity is immediately apparent in areas within the same prefrontal regions, e.g. areas 13 and 14 sharing reciprocal connectivity (Barbas & Pandya, 1989; Carmichael & Price, 1996), and in areas bounded by each other in separate regions, e.g. 9 and 10 (Barbas & Pandya 1989; Petrides & Pandya 2007). Our results affirm this rule principle. Connections between neighboring areas within the same PFC partition (e.g. 9 and 9/46) and across partition boundaries (e.g. 46 and 47) were consistently reconstructed as dense, plausible bundles and classified as Robust True Positives. It is possible that the consistent connectivity of adjacent cortical areas is due to the inherent tendency in tractography to reconstruct spatially proximal pathways (Liptrot et al., 2014). However, we attribute our findings to underlying biology rather than imaging bias for the following reasons. As previously described, these tracts reflect a consistent biological precedence established from the tract-tracing literature. Further, these short-range tracts closely adhere to the surrounding anatomy on an individual level, as these tracts tend to follow the curvature of surrounding gyri and sulci. Lastly, our findings show that proximal regions often displayed true negatives (i/e. areas 11 & 25, 44 & 10). While these regions are not adjacent, this biologically justified absence of streamlines suggests that proximity was not the only condition necessary for streamline reconstruction.

Our tractography also mirrored regional connectivity patterns. Both histology and our tractography findings suggest that lateral prefrontal regions, including the dl-PFC, vl-PFC, and lateral frontal pole, are strongly interconnected (Haber et al., 2022). Beyond these broad patterns, we replicated finer distinctions: for example, histological findings suggest that out of the dl-PFC, areas 8 and 9 most consistently display connectivity to the longer-range ACC, with strong connections to areas 24 and 32 beyond other dl-PFC and ACC regions (Pandya et al., 1981; Barbas & Pandya 1989; Carmichael & Price 1996; Petrides & Pandya 1999; Petrides & Pandya, 2006; Haber et al., 2022). We replicated these findings in our dl-PFC tractography results, where 8 and 9 were the only dl-PFC regions connected to 24 and 32 of the ACC. Histological studies show strong vlPFC to OFC connectivity, particularly between areas 45 and 11, and 47 with the OFC, with no links to area 44 (Carmichael & Price, 1996; Schmahmann & Pandya, 2006; Petrides & Pandya, 2007; Haber et al., 2022). Our tractography results closely mirrored this pattern, identifying connections between areas 45 and 11 and between 47 and the OFC, while confirming the absence of connections involving area 44. This demonstrates that histology-informed tractography can capture both the broad organization and fine-grained distinctions of PFC connectivity.

### Discrepancies with Histology and Methodological Limitations

While our results generally aligned with histological findings, there were notable discrepancies, highlighting inherent limitations of tractography. We observed both underestimation of known connections (FNs) and generation of implausible bundles (FPs). For example, area 32, which histology suggests to be extensively connected across the PFC (Yeterian et al., 2012; Haber et al., 2022), showed predominantly false positives. Similarly, the connections between the OFC and dl-PFC connections (Barbas & Pandya, 1989; Carmichael & Price, 1996; Petrides & Pandya, 1999) were not reconstructed, as well as many tracts from the ACC. This may reflect a fundamental difference between NHP and human SAF connectivity, as the two species can show significant differences in white matter connectivity (Donahue et al., 2018). However, it is more likely that these differences arise from fundamental limitations of diffusion tractography, as there is strong histological precedence for widespread prefrontal connections stemming from the OFC and ACC (Barbas & Pandya, 1989; Carmichael & Price, 1995a, 1995b, 1996; Koch et al., 2002). For instance, it is possible that the lack of streamlines is a result of the lower signal-to-noise ratio of the ACC and OFC (Koch et al., 2002; Clark et al., 2021). As prefrontal SAFs are by nature small, they are especially susceptible to signal-to-noise distortions and signal dropout. This is further compounded by tractography’s inherent bias towards short-range tracts (Liptrot et al., 2014; Donahue et al., 2016; Schilling et al., 2018), as both the OFC and ACC are structurally more isolated than the lateral and rostral regions of the PFC, making the likelihood of reconstructing a tract increasingly difficult compared to more spatially proximal regions of the PFC. Similarly, many medial tracts stemming from the ACC were susceptible to false positive complications largely arising due to contamination from proximal crossing fibers from deep white matter structures. Notably, many of these medial false positive tracts branch off the cingulum or the genu of the corpus callosum. This makes it difficult to parse out short association fibers, as these medial streamlines often are dominated by the clear, dense connections of the CC or cingulum. While these are discrepancies from the histological literature, we do not believe they undermine our primary findings. Rather, they define the conditions and locations where our histology-informed tractography is most challenged. Future refinements may be needed to overcome challenging geometrical or white matter architectures.

### Test-Retest Reliability and Individual Variability

Our findings show that prefrontal SAFs show high test-retest reliability. While wDice was moderate to strong (range .67–.73), it is comparable to other studies on short (Mendoza et al., 2024) and some long range pathways (Boukadi et al., 2018; Zhang et al., 2019a; Schilling et al., 2021b). Bundle adjacency, which is typically a better indicator of reliability than wDice for smaller structures like these (Liu et al., 2024; Mendoza et al., 2024), revealed strong scan-rescan reproducibility. Our results suggest that between scans, these tracts typically only differ on the order of the size of one voxel (range: 1.01–1.35 mm). This meets and exceeds reproducibility of larger, well-established pathways (Schilling et al., 2021a), demonstrating that the overall location and trajectory of these pathways is highly stable within an individual.

Between-subject variability was much greater than within-subject variability, indicating that prefrontal connections are highly individualized. This aligns with prior findings that the PFC shows consistent individual variability (Bürgel et al., 2006; Petrides et al., 2012), which may underlie individual differences in cognition and behavior and provide a baseline for detecting abnormalities in clinical populations.

Taken together, these findings suggest that prefrontal SAFs can be measured with a high degree of reliability. More importantly, they provide a quantitative measurement or feature for individual variability, offering a new avenue for investigating the anatomical basis of human individuality in both health and disease.

### Potential Neuroanatomical Insights

Beyond validating well-established pathways, the synergy between histology and tractography provides a blueprint for extending anatomical knowledge, particularly in cortical regions that are underexplored in the primate literature, such as the frontal pole (area 10). Due to its significant evolutionary expansion in humans relative to non-human primates (Semendeferi et al., 2001), its connective architecture is less extensively mapped histologically (Semendeferi et al., 2001; Petrides & Pandya, 2007; Haber et al., 2022).

Our tractography of the frontal pole, which achieved reliability and accuracy of 100% against the literature, robustly corroborated the sparse existing findings, such as its widespread connectivity and lack of connections to area 44. Crucially, our data extends this foundational knowledge. We reveal that these connections are not monolithic, but are volumetrically large, densely packed, and often organized into distinct sub-bundles.

This demonstrates how this approach can generate novel, detailed hypotheses about the connective architecture of underexplored regions. It provides a reliable in-vivo map that corroborates the available histology while filling in critical gaps. Future work using streamline clustering techniques to delineate these sub-bundles represents a promising next step to probe the functional organization of these newly characterized pathways (Vázquez et al., 2020).

### Considerations and Limitations

There are limitations to this study. First, quality control of reconstructed bundles was performed by a single rater with formal training in neuroanatomy, introducing the potential for rater bias. However, the distinction between tractography classifications in our dataset was typically not ambiguous. The True Positive classifications reflected whether a given connection was observed in more than 50% of subjects, but in practice, the majority of connections were reconstructed in a large majority of subjects (often >80%). Only 8 connections fell into the TP category, whereas the remaining 41 were RTP, indicating that most classifications were far from the decision boundary and unlikely to be affected by rater uncertainty. Consistent with this, connections classified as a True or False Negative generally showed either no streamlines or only a negligible number (e.g., fewer than 20 streamlines), making their classification unambiguous. Lastly, to ensure reproducibility of rating, a second rater conducted quality control of a subsample of 6,000 individual tracts. We then compared rater 2’s classifications to rater 1’s classifications and found 97% agreement between the two raters, suggesting a high degree of consistency. We believe that the stark separation observed between bundle classifications in this dataset as well as our subsample consistency makes it unlikely that rater bias meaningfully influenced our conclusions.

Another potential limitation concerns organizational differences between the NHP and human brain. While the PFC has undergone evolutionary expansion from NHPs to humans, recent studies suggest that the primary differences between the two species are not organizational, as the general cytoarchitectural and connectomic organization of the PFC appears to be remarkably well preserved across species. Rather, recent studies indicate that the principal distinction lies in relative size, with the human PFC being substantially larger than that of NHPs (Petrides et al., 2012; Jbabdi et al., 2013; Donahue et al., 2018; Levy, 2024). This prefrontal expansion is accompanied by increased cortical gyrification (Levy, 2024), motivating our decision to define ROIs using cytoarchitecturally based Brodmann Areas rather than macroscopic regional landmarks, which may be less consistent across species. Indeed, prior work suggests that expansion of the human brain largely maintained the organizational blueprint observed in the rhesus macaque (Petrides et al., 2012; Jbabdi et al., 2013; Sallet et al., 2013; Levy, 2024), and functional coupling studies have failed to identify novel prefrontal regions unique to humans (Sallet et al., 2013). Together, these findings suggest that the organizational properties of short-range PFC connections are likely conserved across species. Nonetheless, future work should employ tractography across species to more directly cross-validate these findings.

It is important to mention that, despite its limitations, non-human primate tract-tracing literature is currently the closest the tractography community has in terms of ground-truth anatomy. While human Klinger dissection studies can elucidate overall organizational characteristics of long-range tracts such as the corpus callosum, corticospinal tract, and superior longitudinal fasciculus with a human brain, they are unable to provide the precise cytoarchitectural cortical terminations for these connections (Martino et al., 2011), let alone the small, complex, and intricate short-fibers of the superficial white matter, although advances in dissection techniques may facilitate this area of research (Dannhoff et al., 2024). Instead, the histological literature is a valuable tool to tractography investigators as it clearly and elaborately lists the terminations of each Brodmann Area, giving us the best current guidance on a-priori tract reconstruction for these small SAFs. The main differences between human and monkey PFC interconnections may lie within the nuances and subdivisions of the interregional connectivity. This means that overall regional connectivity is preserved (i/e., dl-PFC to vl-PFC; area 9 to 10), as this is what both structural and functional studies suggest. It is more possible that the organizational differences in structural connectivity between species lie in nuanced regional subdivisions (i/e., 9l to 8Ad vs 9m to 8BM). In this study, we combined corresponding BA ROI’s from the HCP-MMP1 atlas (i/e, 8A + 8B + 8C into a singular BA 8 ROI). However, tract-tracing literature consistently shows that there are meaningfully different bundles connecting subregions of different Brodmann Areas. For example, areas 8C and 9/46d are connected, and areas 8B and 9/46v are connected in differentiable patterns, rather than reporting the connection between areas 8 and 9/46 (Petrides & Pandya, 1999). While this is not feasible within the scope of this study, this is an important area of future investigation.

Lastly, partial volume effects represent an inherent limitation of diffusion tractography. This issue even affects the accuracy of large, canonical deep white-matter tract reconstruction. It is particularly relevant in the present study, as SAFs of the PFC are volumetrically small. While we leveraged high-resolution diffusion data from the HCP and grounded our analyses in established histological literature, future studies would benefit from replicating these findings using sub-millimeter diffusion imaging to further mitigate partial volume effects.

## Conclusion

This study addresses a long-standing gap in the human connectome: visualizing short-range connections of the human prefrontal cortex. By demonstrating that tractography, when guided by the biological ground truth of NHP tract-tracing, can reconstruct these pathways with high fidelity and precision, we establish a robust framework for mapping an understudied and previously inaccessible component of the human connectome. These connections are anatomically faithful, robust, and highly individualized. Ultimately, this work bridges the gap between invasive animal studies and non-invasive human neuroscience, opening new avenues to investigate the local circuitry underlying complex behavior in health and disease.

## Acknowledgements

This study was funded by NIH T32 EB001628, K01 EB032898, and R01 EB017230.

## Footnotes

1 While the confusion matrix metrics above describe connections (i.e., true positive *connections*, or false positive *connections*), it is critical to also consider the trajectory that tractography follows when classifying accuracy. For this reason, we consider an anatomically implausible tractography trajectory (which is consistent across subjects) to be a false positive connection, regardless of the presence/absence of histology, in line with Maier-Hein et al. 2017.

## References

Amandola, M., Farber, K., Kidambi, R., & Leung (梁海松), H.-C. (2025). Large-Scale High-Resolution Probabilistic Maps of the Human Superior Longitudinal Fasciculus Subdivisions and Their Cortical Terminations. The Journal of Neuroscience, 45(18), e0821242025. 10.1523/JNEUROSCI.0821-24.2025

Avants, B., Epstein, C., Grossman, M., & Gee, J. (2008). Symmetric diffeomorphic image registration with cross-correlation: Evaluating automated labeling of elderly and neurodegenerative brain. Medical Image Analysis, 12(1), 26–41. 10.1016/j.media.2007.06.004

Barbas, H., & Pandya, D. N. (1989). Architecture and intrinsic connections of the prefrontal cortex in the rhesus monkey. The Journal of Comparative Neurology, 286(3), 353–375. 10.1002/cne.902860306

Bookheimer, S. Y., Salat, D. H., Terpstra, M., Ances, B. M., Barch, D. M., Buckner, R. L., Burgess, G. C., Curtiss, S. W., Diaz-Santos, M., Elam, J. S., Fischl, B., Greve, D. N., Hagy, H. A., Harms, M. P., Hatch, O. M., Hedden, T., Hodge, C., Japardi, K. C., Kuhn, T. P., … Yacoub, E. (2019). The Lifespan Human Connectome Project in Aging: An overview. NeuroImage, 185, 335–348. 10.1016/j.neuroimage.2018.10.009

Boukadi, M., Marcotte, K., Bedetti, C., Houde, J.-C., Desautels, A., Deslauriers-Gauthier, S., Chapleau, M., Boré, A., Descoteaux, M., & Brambati, S. M. (2019). Test-Retest Reliability of Diffusion Measures Extracted Along White Matter Language Fiber Bundles Using HARDI-Based Tractography. Frontiers in Neuroscience, 12, 1055. 10.3389/fnins.2018.01055

Bozkurt, B., Yagmurlu, K., Middlebrooks, E. H., Karadag, A., Ovalioglu, T. C., Jagadeesan, B., Sandhu, G., Tanriover, N., & Grande, A. W. (2016). Microsurgical and Tractographic Anatomy of the Supplementary Motor Area Complex in Humans. World Neurosurgery, 95, 99–107. 10.1016/j.wneu.2016.07.072

Brodmann, K. (1909). Vergleichende Lokalisationslehre der Grosshirnrinde in ihren Prinzipien dargestellt auf Grund des Zellenbaues (J. A. Barth, 1909); Brodmann’s Localization in the Cerebral Cortex. Smith Gordon, 1994 [transl. Garey, L.J.].

Bürgel, U., Amunts, K., Hoemke, L., Mohlberg, H., Gilsbach, J. M., & Zilles, K. (2006). White matter fiber tracts of the human brain: Three-dimensional mapping at microscopic resolution, topography and intersubject variability. NeuroImage, 29(4), 1092–1105. 10.1016/j.neuroimage.2005.08.040

Burks, J. D., Boettcher, L. B., Conner, A. K., Glenn, C. A., Bonney, P. A., Baker, C. M., Briggs, R. G., Pittman, N. A., O’Donoghue, D. L., Wu, D. H., & Sughrue, M. E. (2017b). White matter connections of the inferior parietal lobule: A study of surgical anatomy. Brain and Behavior, 7(4), e00640. 10.1002/brb3.640

Carmichael, S. T., & Price, J. L. (1995a). Limbic connections of the orbital and medial prefrontal cortex in macaque monkeys. The Journal of Comparative Neurology, 363(4), 615–641. 10.1002/cne.903630408

Carmichael, S. T., & Price, J. L. (1995b). Sensory and premotor connections of the orbital and medial prefrontal cortex of macaque monkeys. The Journal of Comparative Neurology, 363(4), 642–664. 10.1002/cne.903630409

Carmichael, S. T., & Price, J. L. (1996). Connectional networks within the orbital and medial prefrontal cortex of macaque monkeys. The Journal of Comparative Neurology, 371(2), 179–207. 10.1002/(SICI)1096-9861(19960722)371:2%253C179::AID-CNE1%253E3.0.CO;2-%2523

Catani, M., Dell’acqua, F., Vergani, F., Malik, F., Hodge, H., Roy, P., Valabregue, R., & Thiebaut de Schotten, M. (2012). Short frontal lobe connections of the human brain. Cortex; a Journal Devoted to the Study of the Nervous System and Behavior, 48(2), 273–291. 10.1016/j.cortex.2011.12.001

Cavada, C., Compañy, T., Tejedor, J., Cruz-Rizzolo, R. J., & Reinoso-Suárez, F. (2000). The anatomical connections of the macaque monkey orbitofrontal cortex. A review. Cerebral Cortex (New York, N.Y.: 1991), 10(3), 220–242. 10.1093/cercor/10.3.220

Clark, I. A., Callaghan, M. F., Weiskopf, N., Maguire, E. A., & Mohammadi, S. (2021). Reducing Susceptibility Distortion Related Image Blurring in Diffusion MRI EPI Data. Frontiers in Neuroscience, 15, 706473. 10.3389/fnins.2021.706473

Cousineau, M., Jodoin, P.-M., Morency, F. C., Rozanski, V., Grand’Maison, M., Bedell, B. J., & Descoteaux, M. (2017). A test-retest study on Parkinson’s PPMI dataset yields statistically significant white matter fascicles. NeuroImage. Clinical, 16, 222–233. 10.1016/j.nicl.2017.07.020

Dannhoff, G., Morichon, A., Smirnov, M., Barantin, L., Destrieux, C., & Maldonado, I. L. (2024). Direct Inside-Out Observation of Superficial White Matter Fasciculi in the Human Brain. Brain Connectivity, 14(2), 107–121. 10.1089/brain.2023.0050

De Schotten, M. T., Dell’Acqua, F., Forkel, S. J., Simmons, A., Vergani, F., Murphy, D. G. M., & Catani, M. (2011). A lateralized brain network for visuospatial attention. Nature Neuroscience, 14(10), 1245–1246. 10.1038/nn.2905

Dice, L. R. (1945). Measures of the Amount of Ecologic Association Between Species. Ecology, 26(3), 297–302. 10.2307/1932409

Donahue, C. J., Glasser, M. F., Preuss, T. M., Rilling, J. K., & Van Essen, D. C. (2018a). Quantitative assessment of prefrontal cortex in humans relative to nonhuman primates. Proceedings of the National Academy of Sciences, 115(22). 10.1073/pnas.1721653115

Donahue, C. J., Sotiropoulos, S. N., Jbabdi, S., Hernandez-Fernandez, M., Behrens, T. E., Dyrby, T. B., Coalson, T., Kennedy, H., Knoblauch, K., Van Essen, D. C., & Glasser, M. F. (2016). Using Diffusion Tractography to Predict Cortical Connection Strength and Distance: A Quantitative Comparison with Tracers in the Monkey. The Journal of Neuroscience, 36(25), 6758–6770. 10.1523/JNEUROSCI.0493-16.2016

Dyrby, T. B., Søgaard, L. V., Parker, G. J., Alexander, D. C., Lind, N. M., Baaré, W. F. C., Hay-Schmidt, A., Eriksen, N., Pakkenberg, B., Paulson, O. B., & Jelsing, J. (2007). Validation of in vitro probabilistic tractography. NeuroImage, 37(4), 1267–1277. 10.1016/j.neuroimage.2007.06.022

Fischl, B. (2012). FreeSurfer. NeuroImage, 62(2), 774–781. 10.1016/j.neuroimage.2012.01.021

Fonov, V., Evans, A. C., Botteron, K., Almli, C. R., McKinstry, R. C., Collins, D. L., & Brain Development Cooperative Group. (2011). Unbiased average age-appropriate atlases for pediatric studies. NeuroImage, 54(1), 313–327. 10.1016/j.neuroimage.2010.07.033

Frey, S., Mackey, S., & Petrides, M. (2014). Cortico-cortical connections of areas 44 and 45B in the macaque monkey. Brain and Language, 131, 36–55. 10.1016/j.bandl.2013.05.005

Funahashi, S., Chafee, M. V., & Goldman-Rakic, P. S. (1993). Prefrontal neuronal activity in rhesus monkeys performing a delayed anti-saccade task. Nature, 365(6448), 753–756. 10.1038/365753a0

Gerbella, M., Belmalih, A., Borra, E., Rozzi, S., & Luppino, G. (2010). Cortical connections of the macaque caudal ventrolateral prefrontal areas 45A and 45B. Cerebral Cortex (New York, N.Y.: 1991), 20(1), 141–168. 10.1093/cercor/bhp087

Girard, G., Caminiti, R., Battaglia-Mayer, A., St-Onge, E., Ambrosen, K. S., Eskildsen, S. F., Krug, K., Dyrby, T. B., Descoteaux, M., Thiran, J.-P., & Innocenti, G. M. (2020b). On the cortical connectivity in the macaque brain: A comparison of diffusion tractography and histological tracing data. NeuroImage, 221, 117201. 10.1016/j.neuroimage.2020.117201

Glasser, M. F., Coalson, T. S., Robinson, E. C., Hacker, C. D., Harwell, J., Yacoub, E., Ugurbil, K., Andersson, J., Beckmann, C. F., Jenkinson, M., Smith, S. M., & Van Essen, D. C. (2016). A multi-modal parcellation of human cerebral cortex. Nature, 536(7615), 171–178. 10.1038/nature18933

Glasser, M. F., Sotiropoulos, S. N., Wilson, J. A., Coalson, T. S., Fischl, B., Andersson, J. L., Xu, J., Jbabdi, S., Webster, M., Polimeni, J. R., Van Essen, D. C., Jenkinson, M., & WU-Minn HCP Consortium. (2013). The minimal preprocessing pipelines for the Human Connectome Project. NeuroImage, 80, 105–124. 10.1016/j.neuroimage.2013.04.127

Guevara, M., Guevara, P., Román, C., & Mangin, J.-F. (2020). Superficial white matter: A review on the dMRI analysis methods and applications. NeuroImage, 212, 116673. 10.1016/j.neuroimage.2020.116673

Guevara, M., Román, C., Houenou, J., Duclap, D., Poupon, C., Mangin, J. F., & Guevara, P. (2017). Reproducibility of superficial white matter tracts using diffusion-weighted imaging tractography. NeuroImage, 147, 703–725. 10.1016/j.neuroimage.2016.11.066

Guevara, M., Sun, Z.-Y., Guevara, P., Rivière, D., Grigis, A., Poupon, C., & Mangin, J.-F. (2022). Disentangling the variability of the superficial white matter organization using regional-tractogram-based population stratification. NeuroImage, 255, 119197. 10.1016/j.neuroimage.2022.119197

Haber, S. N., Liu, H., Seidlitz, J., & Bullmore, E. (2022). Prefrontal connectomics: From anatomy to human imaging. Neuropsychopharmacology: Official Publication of the American College of Neuropsychopharmacology, 47(1), 20–40. 10.1038/s41386-021-01156-6

Jbabdi, S., Lehman, J. F., Haber, S. N., & Behrens, T. E. (2013). Human and monkey ventral prefrontal fibers use the same organizational principles to reach their targets: Tracing versus tractography. The Journal of Neuroscience: The Official Journal of the Society for Neuroscience, 33(7), 3190–3201. 10.1523/JNEUROSCI.2457-12.2013

Jeurissen, B., Tournier, J.-D., Dhollander, T., Connelly, A., & Sijbers, J. (2014). Multi-tissue constrained spherical deconvolution for improved analysis of multi-shell diffusion MRI data. NeuroImage, 103, 411–426. 10.1016/j.neuroimage.2014.07.061

Knösche, T. R., Anwander, A., Liptrot, M., & Dyrby, T. B. (2015). Validation of tractography: Comparison with manganese tracing. Human Brain Mapping, 36(10), 4116–4134. 10.1002/hbm.22902

Koch, M. A., Glauche, V., Finsterbusch, J., Nolte, U. G., Frahm, J., Weiller, C., & Büchel, C. (2002). Distortion-free Diffusion Tensor Imaging of Cranial Nerves and of Inferior Temporal and Orbitofrontal White Matter. NeuroImage, 17(1), 497–506. 10.1006/nimg.2002.1171

Levy, R. (2024). The prefrontal cortex: From monkey to man. Brain: A Journal of Neurology, 147(3), 794–815. 10.1093/brain/awad389

Levy, R., & Goldman-Rakic, P. S. (2000). Segregation of working memory functions within the dorsolateral prefrontal cortex. Experimental Brain Research, 133(1), 23–32. 10.1007/s002210000397

Liptrot, M. G., Sidaros, K., & Dyrby, T. B. (2014). Addressing the Path-Length-Dependency Confound in White Matter Tract Segmentation. PLoS ONE, 9(5), e96247. 10.1371/journal.pone.0096247

Liu, B., Dolz, J., Galdran, A., Kobbi, R., & Ben Ayed, I. (2024). Do we really need dice? The hidden region-size biases of segmentation losses. Medical Image Analysis, 91, 103015. 10.1016/j.media.2023.103015

Maier-Hein, K. H., Neher, P. F., Houde, J.-C., Côté, M.-A., Garyfallidis, E., Zhong, J., Chamberland, M., Yeh, F.-C., Lin, Y.-C., Ji, Q., Reddick, W. E., Glass, J. O., Chen, D. Q., Feng, Y., Gao, C., Wu, Y., Ma, J., He, R., Li, Q., … Descoteaux, M. (2017). The challenge of mapping the human connectome based on diffusion tractography. Nature Communications, 8(1), 1349. 10.1038/s41467-017-01285-x

Marek, S., & Dosenbach, N. U. F. (2018). The frontoparietal network: Function, electrophysiology, and importance of individual precision mapping. Dialogues in Clinical Neuroscience, 20(2), 133–140. 10.31887/DCNS.2018.20.2/smarek

Martino, J., De Witt Hamer, P. C., Vergani, F., Brogna, C., de Lucas, E. M., Vázquez-Barquero, A., García-Porrero, J. A., & Duffau, H. (2011). Cortex-sparing fiber dissection: An improved method for the study of white matter anatomy in the human brain. Journal of Anatomy, 219(4), 531–541. 10.1111/j.1469-7580.2011.01414.x

Mendoza, C., Román, C., Mangin, J.-F., Hernández, C., & Guevara, P. (2024). Short fiber bundle filtering and test-retest reproducibility of the Superficial White Matter. Frontiers in Neuroscience, 18, 1394681. 10.3389/fnins.2024.1394681

Morecraft, R. J., Stilwell-Morecraft, K. S., Cipolloni, P. B., Ge, J., McNeal, D. W., & Pandya, D. N. (2012). Cytoarchitecture and cortical connections of the anterior cingulate and adjacent somatomotor fields in the rhesus monkey. Brain Research Bulletin, 87(4–5), 457–497. 10.1016/j.brainresbull.2011.12.005

Movahedian Attar, F., Kirilina, E., Chaimow, D., Haenelt, D., Schneider, C., Edwards, L. J., Pine, K. J., Jäger, C., Reimann, K., Pohlmann, A., Periquito, J., Streubel, T., Trampel, R., Mohammadi, S., Niendorf, T., Morawski, M., & Weiskopf, N. (2025). Short association fibres form topographic sheets in the human V1–V2 processing stream. Imaging Neuroscience, 3, imag_a_00498. 10.1162/imag_a_00498

Nazeri, A., Chakravarty, M. M., Rajji, T. K., Felsky, D., Rotenberg, D. J., Mason, M., Xu, L. N., Lobaugh, N. J., Mulsant, B. H., & Voineskos, A. N. (2015). Superficial white matter as a novel substrate of age-related cognitive decline. Neurobiology of Aging, 36(6), 2094–2106. 10.1016/j.neurobiolaging.2015.02.022

Pandya, D. N., Van Hoesen, G. W., & Mesulam, M. M. (1981). Efferent connections of the cingulate gyrus in the rhesus monkey. Experimental Brain Research, 42(3–4), 319–330. 10.1007/BF00237497

Petrides, M., & Pandya, D. N. (1999). Dorsolateral prefrontal cortex: Comparative cytoarchitectonic analysis in the human and the macaque brain and corticocortical connection patterns. The European Journal of Neuroscience, 11(3), 1011–1036. 10.1046/j.1460-9568.1999.00518.x

Petrides, M., & Pandya, D. N. (2002). Comparative cytoarchitectonic analysis of the human and the macaque ventrolateral prefrontal cortex and corticocortical connection patterns in the monkey. European Journal of Neuroscience, 16(2), 291–310. 10.1046/j.1460-9568.2001.02090.x

Petrides, M., & Pandya, D. N. (2006). Efferent association pathways originating in the caudal prefrontal cortex in the macaque monkey. The Journal of Comparative Neurology, 498(2), 227–251. 10.1002/cne.21048

Petrides, M., & Pandya, D. N. (2007). Efferent Association Pathways from the Rostral Prefrontal Cortex in the Macaque Monkey. The Journal of Neuroscience, 27(43), 11573–11586. 10.1523/JNEUROSCI.2419-07.2007

Petrides, M., Tomaiuolo, F., Yeterian, E. H., & Pandya, D. N. (2012). The prefrontal cortex: Comparative architectonic organization in the human and the macaque monkey brains. Cortex; a Journal Devoted to the Study of the Nervous System and Behavior, 48(1), 46–57. 10.1016/j.cortex.2011.07.002

Pietrasik, W., Cribben, I., Olsen, F., & Malykhin, N. (2023). Diffusion tensor imaging of superficial prefrontal white matter in healthy aging. Brain Research, 1799, 148152. 10.1016/j.brainres.2022.148152

Preuss, T. M., & Goldman-Rakic, P. S. (1989). Connections of the ventral granular frontal cortex of macaques with perisylvian premotor and somatosensory areas: Anatomical evidence for somatic representation in primate frontal association cortex. The Journal of Comparative Neurology, 282(2), 293–316. 10.1002/cne.902820210

Renauld, E., Boré, A., Poirier, C., Valcourt-Caron, A., Karan, P., Théberge, A., Théaud, G., Edde, M., Poulin, P., Girard, G., Houde, J.-C., Gagnon, A., St-Onge, E., Little, G., Legarreta, J. H., Thoumyre, S., Grenier, G., El Yamani, Z., Ocampo Pineda, M., … Descoteaux, M. (2026). Tractography analysis with the scilpy toolbox. Aperture Neuro, 6. 10.52294/001c.154022

Rheault, F., De Benedictis, A., Daducci, A., Maffei, C., Tax, C. M. W., Romascano, D., Caverzasi, E., Morency, F. C., Corrivetti, F., Pestilli, F., Girard, G., Theaud, G., Zemmoura, I., Hau, J., Glavin, K., Jordan, K. M., Pomiecko, K., Chamberland, M., Barakovic, M., … Descoteaux, M. (2020). Tractostorm: The what, why, and how of tractography dissection reproducibility. Human Brain Mapping, 41(7), 1859–1874. 10.1002/hbm.24917

Rheault, F., Houde, J.-C., Goyette, N., Morency, F. C., & Descoteaux, M. (2016). MI-Brain, a software to handle tractograms and perform interactive virtual dissection. Diffusion Study Group Workshop. International Society for Magnetic Resonance in Medicine.

Rolls, E. T., Rauschecker, J. P., Deco, G., Huang, C.-C., & Feng, J. (2023). Auditory cortical connectivity in humans. Cerebral Cortex, 33(10), 6207–6227. 10.1093/cercor/bhac496

Román, C., Hernández, C., Figueroa, M., Houenou, J., Poupon, C., Mangin, J.-F., & Guevara, P. (2022). Superficial white matter bundle atlas based on hierarchical fiber clustering over probabilistic tractography data. NeuroImage, 262, 119550. 10.1016/j.neuroimage.2022.119550

Sallet, J., Mars, R. B., Noonan, M. P., Neubert, F.-X., Jbabdi, S., O’Reilly, J. X., Filippini, N., Thomas, A. G., & Rushworth, M. F. (2013). The Organization of Dorsal Frontal Cortex in Humans and Macaques. Journal of Neuroscience, 33(30), 12255–12274. 10.1523/JNEUROSCI.5108-12.2013

Schilling, K. G., Archer, D., Yeh, F.-C., Rheault, F., Cai, L. Y., Shafer, A., Resnick, S. M., Hohman, T., Jefferson, A., Anderson, A. W., Kang, H., & Landman, B. A. (2023). Short superficial white matter and aging: A longitudinal multi-site study of 1293 subjects and 2711 sessions. Aging Brain, 3, 100067. 10.1016/j.nbas.2023.100067

Schilling, K. G., Nath, V., Hansen, C., Parvathaneni, P., Blaber, J., Gao, Y., Neher, P., Aydogan, D. B., Shi, Y., Ocampo-Pineda, M., Schiavi, S., Daducci, A., Girard, G., Barakovic, M., Rafael-Patino, J., Romascano, D., Rensonnet, G., Pizzolato, M., Bates, A., … Landman, B. A. (2019). Limits to anatomical accuracy of diffusion tractography using modern approaches. NeuroImage, 185, 1–11. 10.1016/j.neuroimage.2018.10.029

Schilling, K. G., Petit, L., Rheault, F., Remedios, S., Pierpaoli, C., Anderson, A. W., Landman, B. A., & Descoteaux, M. (2020). Brain connections derived from diffusion MRI tractography can be highly anatomically accurate—If we know where white matter pathways start, where they end, and where they do not go. Brain Structure and Function, 225(8), 2387–2402. 10.1007/s00429-020-02129-z

Schilling, K. G., Rheault, F., Petit, L., Hansen, C. B., Nath, V., Yeh, F.-C., Girard, G., Barakovic, M., Rafael-Patino, J., Yu, T., Fischi-Gomez, E., Pizzolato, M., Ocampo-Pineda, M., Schiavi, S., Canales-Rodríguez, E. J., Daducci, A., Granziera, C., Innocenti, G., Thiran, J.-P., … Descoteaux, M. (2021a). Tractography dissection variability: What happens when 42 groups dissect 14 white matter bundles on the same dataset? NeuroImage, 243, 118502. 10.1016/j.neuroimage.2021.118502

Schilling, K. G., Tax, C. M. W., Rheault, F., Hansen, C., Yang, Q., Yeh, F.-C., Cai, L., Anderson, A. W., & Landman, B. A. (2021b). Fiber tractography bundle segmentation depends on scanner effects, vendor effects, acquisition resolution, diffusion sampling scheme, diffusion sensitization, and bundle segmentation workflow. NeuroImage, 242, 118451. 10.1016/j.neuroimage.2021.118451

Schilling, K., Gao, Y., Janve, V., Stepniewska, I., Landman, B. A., & Anderson, A. W. (2018). Confirmation of a gyral bias in diffusion MRI fiber tractography. Human Brain Mapping, 39(3), 1449–1466. 10.1002/hbm.23936

Schilling, K., Zhang, F., Román, C., O’Donnell, L. J., & Guevara, P. (2025). Short association fiber tractography: Key insights and surprising facts. Brain Structure and Function, 230(6), 97. 10.1007/s00429-025-02966-w

Schmahmann, J. D., & Pandya, D. N. (2006). Fiber Pathways of the Brain (1st ed.). Oxford University Press New York. 10.1093/acprof:oso/9780195104233.001.0001

Semendeferi, K., Armstrong, E., Schleicher, A., Zilles, K., & Van Hoesen, G. W. (2001). Prefrontal cortex in humans and apes: A comparative study of area 10. American Journal of Physical Anthropology, 114(3), 224–241. 10.1002/1096-8644(200103)114:3%253C224::AID-AJPA1022%253E3.0.CO;2-I

Shukla, D. K., Keehn, B., Smylie, D. M., & Müller, R.-A. (2011). Microstructural abnormalities of short-distance white matter tracts in autism spectrum disorder. Neuropsychologia, 49(5), 1378–1382. 10.1016/j.neuropsychologia.2011.02.022

Thomas, C., Ye, F. Q., Irfanoglu, M. O., Modi, P., Saleem, K. S., Leopold, D. A., & Pierpaoli, C. (2014). Anatomical accuracy of brain connections derived from diffusion MRI tractography is inherently limited. Proceedings of the National Academy of Sciences of the United States of America, 111(46), 16574–16579. 10.1073/pnas.1405672111

Tournier, J.-D., Smith, R., Raffelt, D., Tabbara, R., Dhollander, T., Pietsch, M., Christiaens, D., Jeurissen, B., Yeh, C.-H., & Connelly, A. (2019). MRtrix3: A fast, flexible and open software framework for medical image processing and visualisation. NeuroImage, 202, 116137. 10.1016/j.neuroimage.2019.116137

Van Dyken, P. C., Khan, A. R., & Palaniyappan, L. (2024). Imaging of the superficial white matter in health and disease. Imaging Neuroscience, 2, 1–35. 10.1162/imag_a_00221

Vavassori, L., Rheault, F., Avesani, P., De Benedictis, A., Corsini, F., Annicchiarico, L., Zigiotto, L., Rozzanigo, U., Barbareschi, M., Petit, L., & Sarubbo, S. (2025). Anatomical insights into the superior longitudinal system from integrative in- vivo and ex-vivo mapping. Communications Biology, 8(1), 1328. 10.1038/s42003-025-08774-6

Vázquez, A., López-López, N., Houenou, J., Poupon, C., Mangin, J.-F., Ladra, S., & Guevara, P. (2020). Automatic group-wise whole-brain short association fiber bundle labeling based on clustering and cortical surface information. Biomedical Engineering Online, 19(1), 42. 10.1186/s12938-020-00786-z

Wasserthal, J., Neher, P., & Maier-Hein, K. H. (2018). TractSeg—Fast and accurate white matter tract segmentation. NeuroImage, 183, 239–253. 10.1016/j.neuroimage.2018.07.070

Wu, Y., Sun, D., Wang, Yong, Wang, Yunjie, & Wang, Yibao. (2016). Tracing short connections of the temporo-parieto-occipital region in the human brain using diffusion spectrum imaging and fiber dissection. Brain Research, 1646, 152–159. 10.1016/j.brainres.2016.05.046

Yeh, F.-C., Panesar, S., Fernandes, D., Meola, A., Yoshino, M., Fernandez-Miranda, J. C., Vettel, J. M., & Verstynen, T. (2018). Population-averaged atlas of the macroscale human structural connectome and its network topology. NeuroImage, 178, 57–68. 10.1016/j.neuroimage.2018.05.027

Yeterian, E. H., Pandya, D. N., Tomaiuolo, F., & Petrides, M. (2012). The cortical connectivity of the prefrontal cortex in the monkey brain. Cortex; a Journal Devoted to the Study of the Nervous System and Behavior, 48(1), 58–81. 10.1016/j.cortex.2011.03.004

Zhang, F., Wu, Y., Norton, I., Rathi, Y., Golby, A. J., & O’Donnell, L. J. (2019a). Test-retest reproducibility of white matter parcellation using diffusion MRI tractography fiber clustering. Human Brain Mapping, 40(10), 3041–3057. 10.1002/hbm.24579

Zhang, F., Wu, Y., Norton, I., Rigolo, L., Rathi, Y., Makris, N., & O’Donnell, L. J. (2018). An anatomically curated fiber clustering white matter atlas for consistent white matter tract parcellation across the lifespan. NeuroImage, 179, 429–447. 10.1016/j.neuroimage.2018.06.027

Zhang, F., Wu, Y., Norton, I., Rigolo, L., Rathi, Y., Makris, N., & O’Donnell, L. J. (2019b). O’Donnell Research Group (ORG) Fiber Clustering White Matter Atlas (Version v1.4) [Dataset]. Zenodo. 10.5281/ZENODO.2648283

