## Supplementary Materials for "Bridging Histology and Tractography: First In-Vivo Visualization of Short-Range Prefrontal Connections Informed by Primate Tract-Tracing"

**Supplementary Figures and Tables**

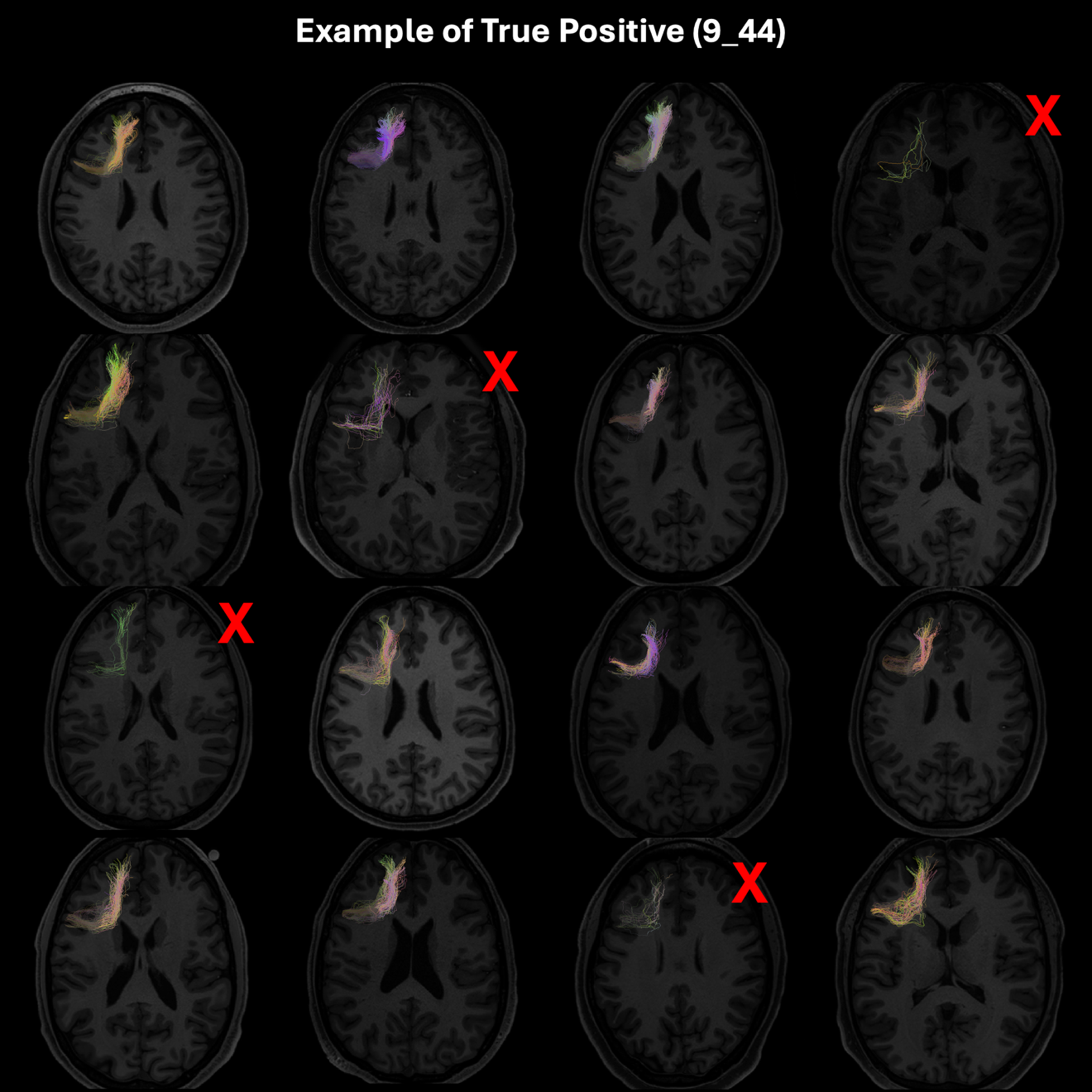

*Supplementary Figure 1 -* Example of bundle classified as a True Positive (TP). TP bundles are anatomically plausible bundles which occur in >50% of the population. All bundles shown are modeled between areas 9 and 44. Red X’s correspond to bundles that did not fail visual inspection. In this case, bundles marked with an X failed due to sparse and random trajectories.

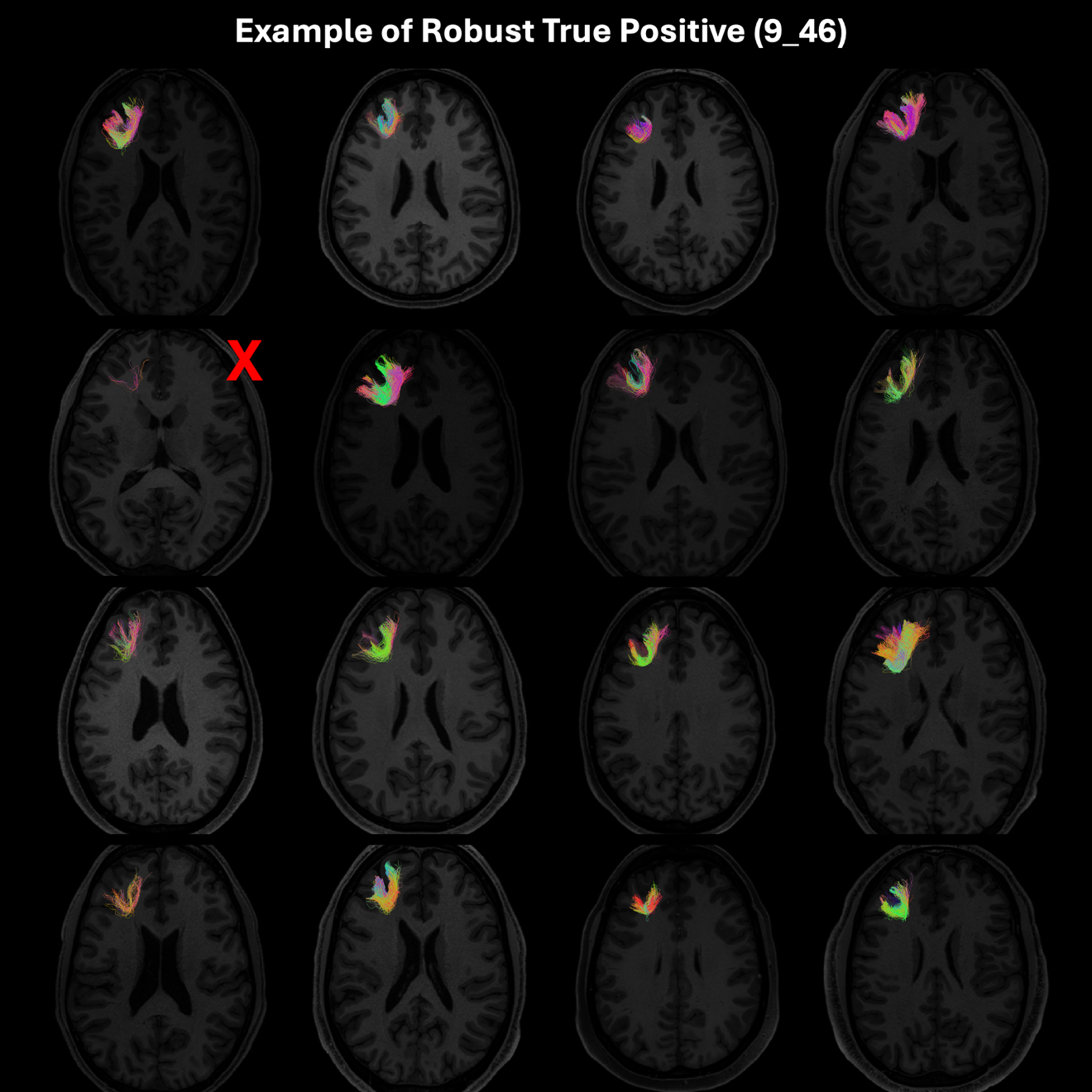

*Supplementary Figure 2* - Example of bundle classified as a Robust True Positive (RTP). RTP bundles are anatomically plausible bundles which occur in >80% of the population. All bundles shown are modeled between areas 9 and 46. Red X’s correspond to bundles that did not fail visual inspection. In this case, the bundle marked with an X failed due to sparse trajectories.

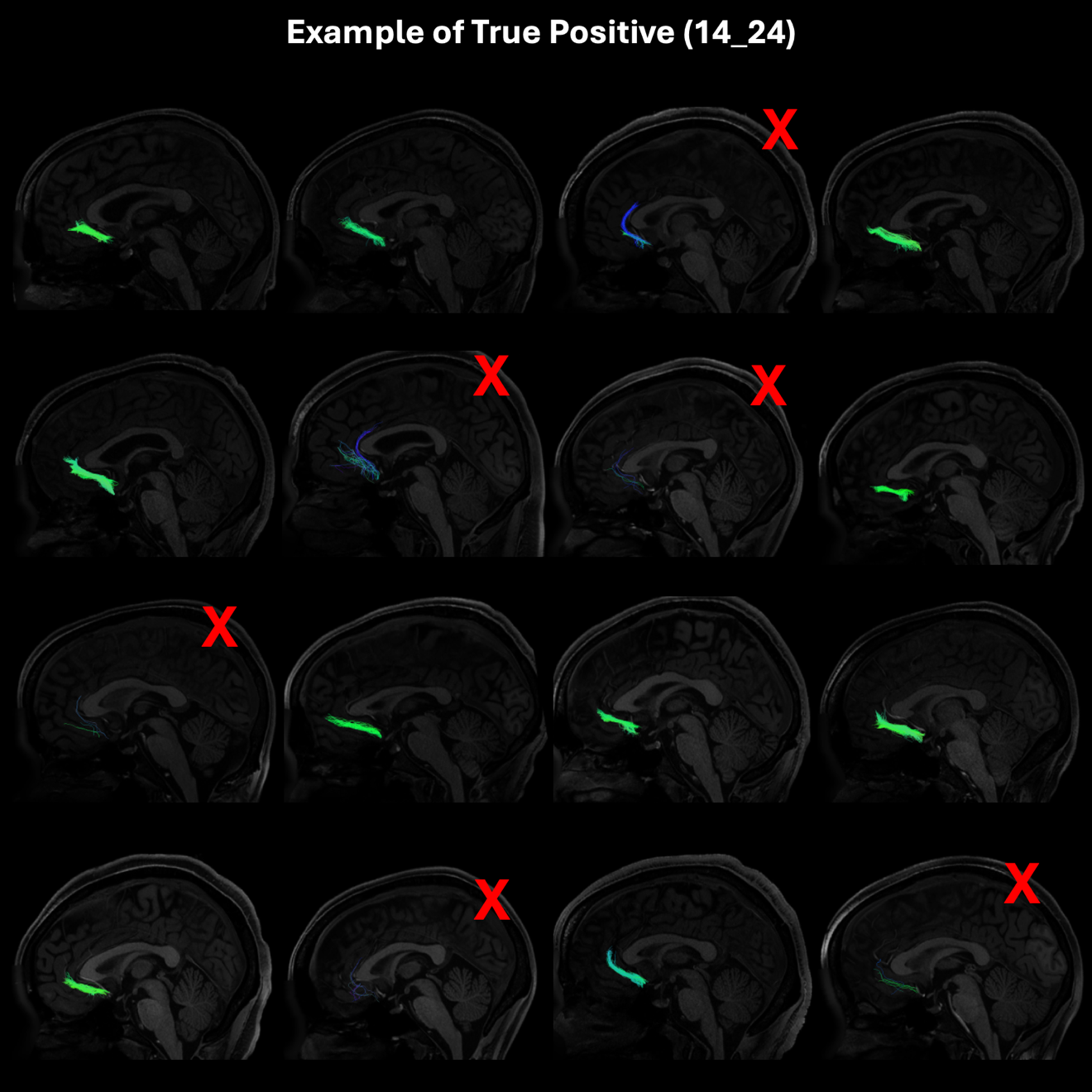

*Supplementary Figure 3* - Additional example of a bundle classified as a True Positive (TP).  All bundles shown are modeled between areas 14 and 24. Red X’s correspond to bundles that did not fail visual inspection. In this case, the bundle marked with an X failed due to sparse trajectories, and trajectories confounded by crossing fibers.

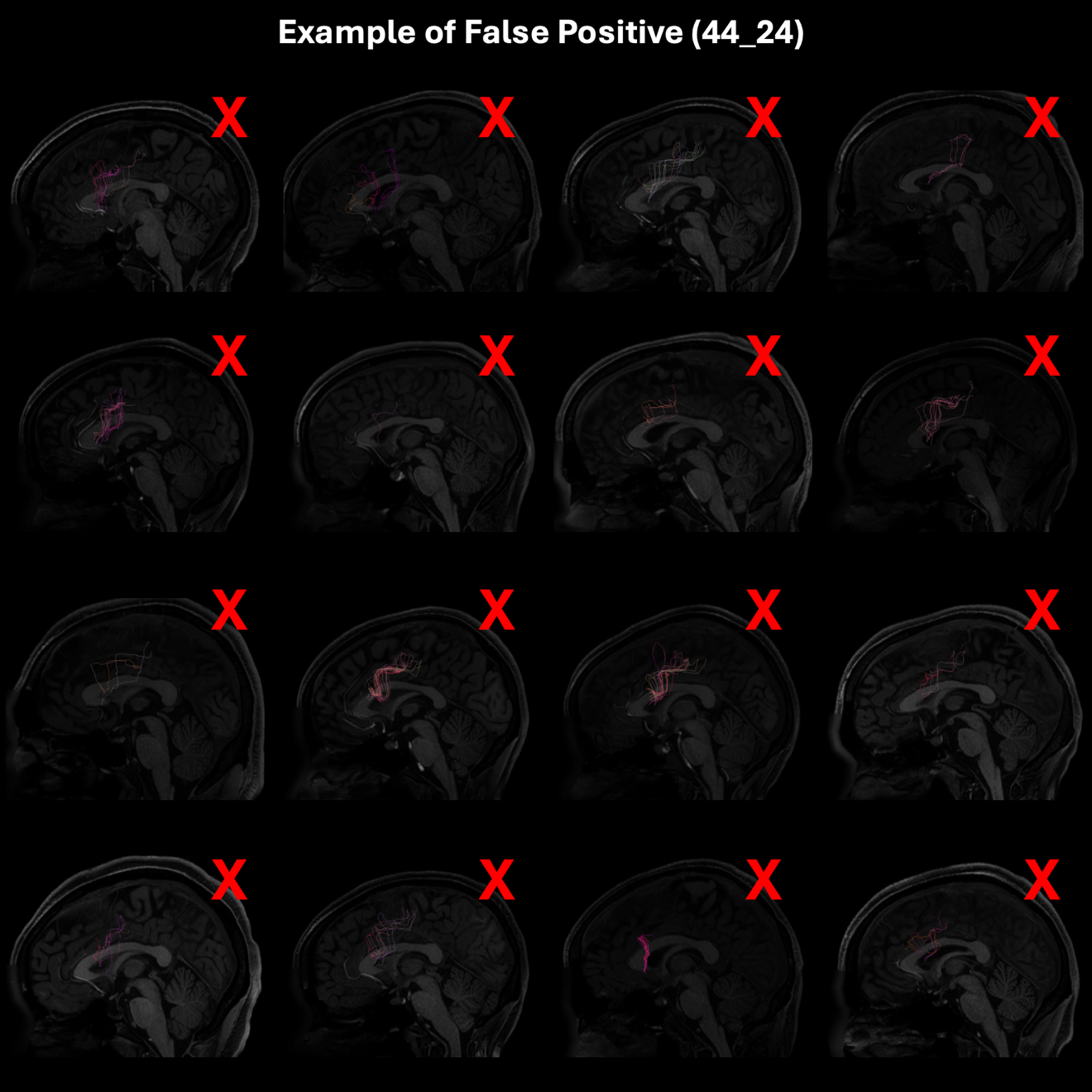

*Supplementary Figure 4* -  Example of bundle classified as a False Positive (FP). FP bundles are anatomically implausible bundles which occur in >50% of the population, or have no histological precedence. All bundles shown are modeled between areas 44 and 24. Red X’s correspond to bundles that did not fail visual inspection. In this case, all bundles failed to sparse, anatomically plausible streamlines.

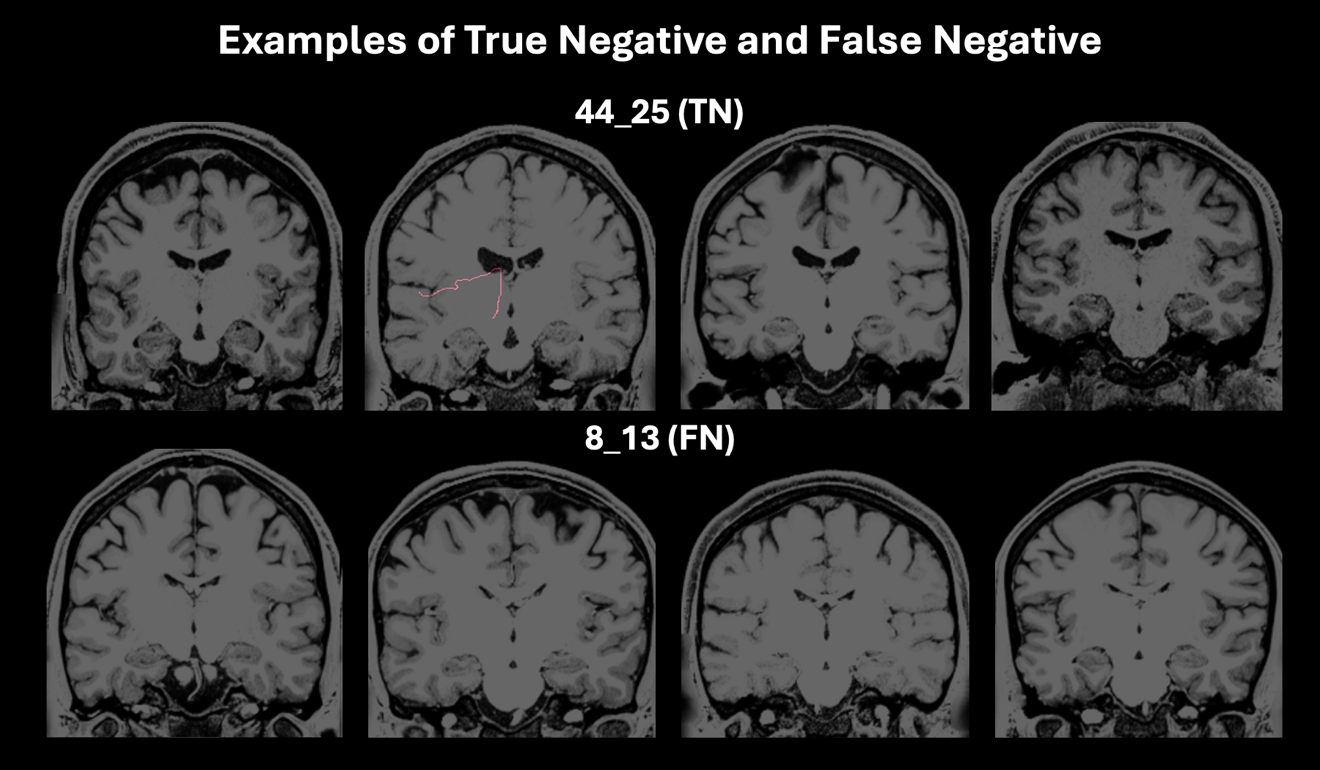

*Supplementary Figure 5* - Examples of bundles classified as True Negatives (TN) and False Negatives (FN). TN bundles generate no streamlines and do not have histological precedence, whereas FN bundles generate no streamlines and do histological precedence.

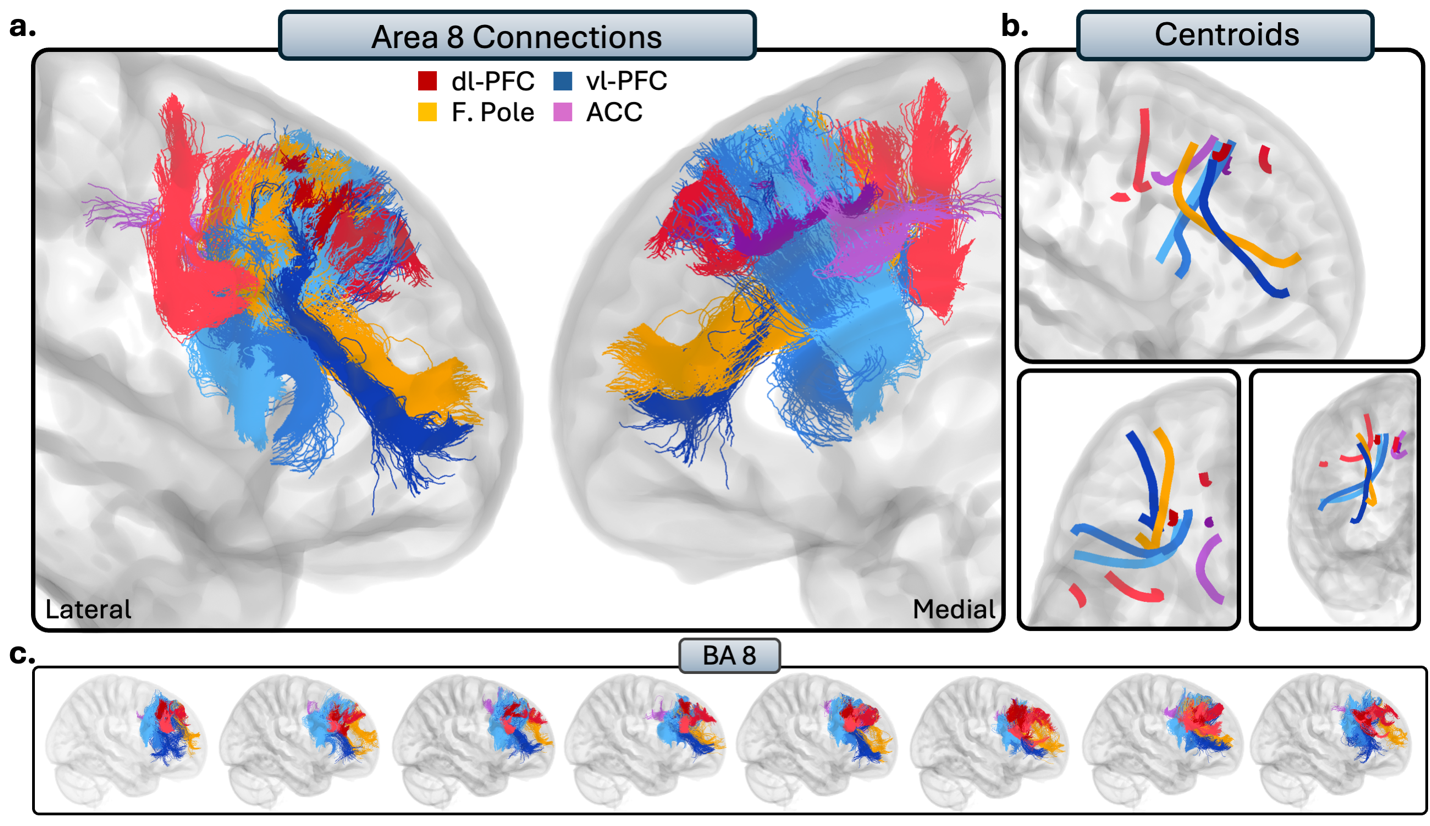

*Supplementary Figure 6 -* As the connections of area 8 are volumetrically large, we separated its visualization from the dl-PFC (Figure 3). a) The connections of area 8 to the rest of the PFC. Red = tracts to the dl-PFC, blue = vl-PFC, orange = frontal pole, and purple = ACC. b.) Overall trajectories of the short fibers. Top: Sagittal view, left: axial view, right: coronal view. c.) Individual subject variability for each prefrontal region.

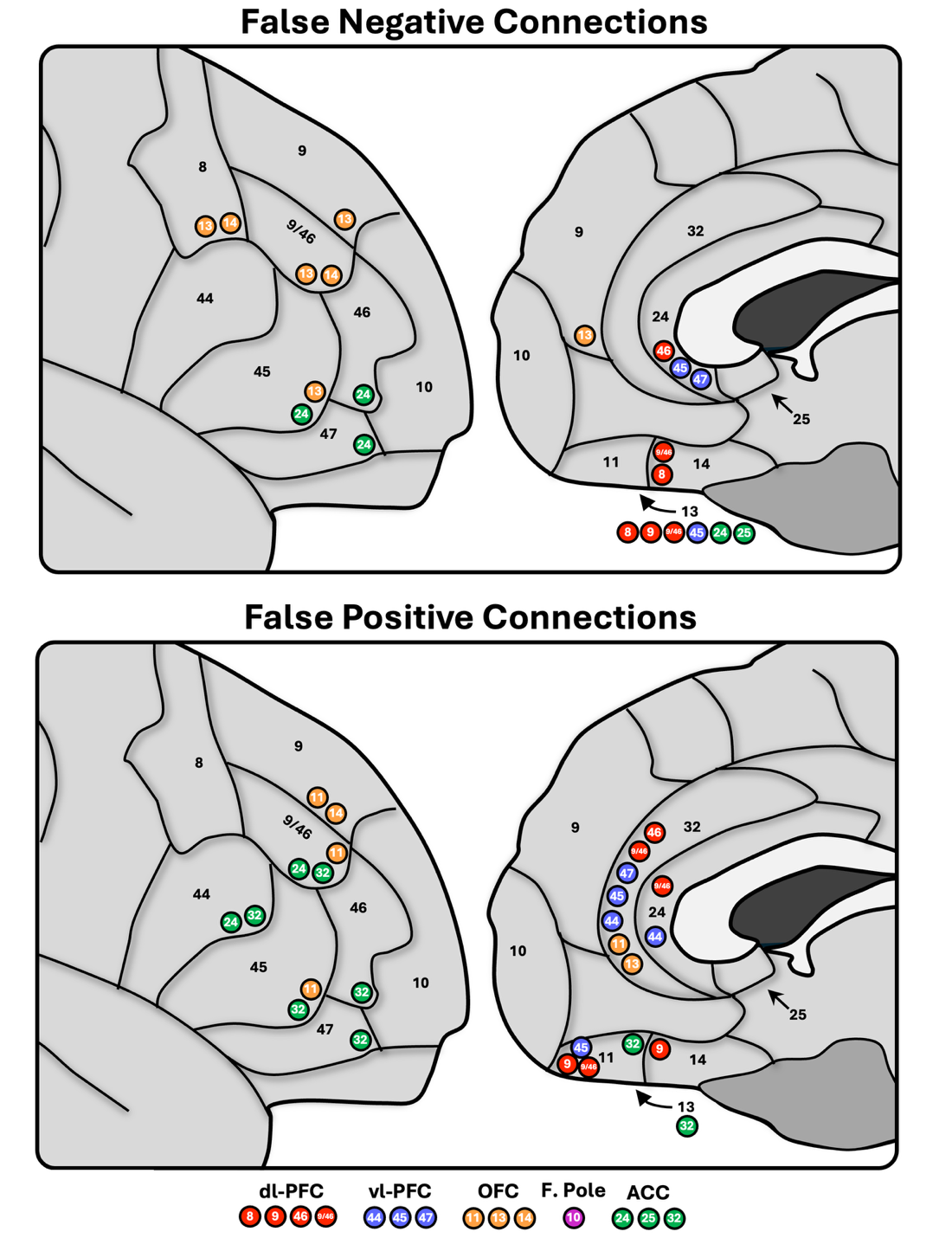

*Supplementary Figure 7* - Schematic overview of the False Negative (FN) and False Positive (FP) tractography results. These are the prefrontal regions where our tractography differed the most from histological findings. Notably, most of these were present in the OFC and ACC.

*Supplementary Table 1* - Maximum Tract Lengths

| **Tract** | ***M* Tract Length (mm)** | **SD** | **Chosen Maximum Length (*M* + 1.5 *SD*)** |
| --- | --- | --- | --- |
| 8_10 | 115.6 | 9.3 | 130 |
| 8_24 | 87.2 | 17.2 | 113 |
| 8_32 | 38.9 | 7.8 | 51 |
| 8_44 | 106.1 | 10.6 | 122 |
| 8_45 | 105.3 | 12.2 | 124 |
| 8_46 | 57.8 | 14.0 | 79 |
| 8_47 | 116.1 | 11.6 | 133 |
| 8_9 | 43.1 | 8.9 | 56 |
| 8_9/46 | 60.6 | 12.8 | 80 |
| 9_10 | 44.4 | 7.5 | 56 |
| 9_24 | 40.3 | 13.5 | 61 |
| 9_32 | 37.8 | 6.5 | 47 |
| 9_44 | 112.2 | 9.7 | 127 |
| 9_45 | 115.8 | 11.0 | 132 |
| 9_46 | 51.9 | 10.0 | 67 |
| 9_47 | 88.4 | 18.3 | 116 |
| 9_9/46 | 53.6 | 11.6 | 71 |
| 9/46_10 | 51.7 | 10.3 | 67 |
| 9/46_11 | 62.9 | 9.5 | 77 |
| 9/46_44 | 92.9 | 14.5 | 115 |
| 9/46_45 | 88.1 | 14.0 | 109 |
| 9/46_47 | 52.2 | 10.5 | 68 |
| 46_10 | 70.4 | 14.8 | 93 |
| 46_32 | 94.4 | 12.1 | 113 |
| 46_44 | 101.9 | 18.2 | 129 |
| 46_45 | 98.3 | 14.0 | 119 |
| 46_47 | 69.1 | 9.5 | 83 |
| 46_9/46 | 53.1 | 10.3 | 69 |
| 44_32 | 100.8 | 10.7 | 117 |
| 44_45 | 49.7 | 10.8 | 66 |
| 45_10 | 100.0 | 14.5 | 122 |
| 45_11 | 98.3 | 13.7 | 119 |
| 45_32 | 105.0 | 9.1 | 119 |
| 47_10 | 63.6 | 17.4 | 90 |
| 47_11 | 47.5 | 11.9 | 65 |
| 47_13 | 39.2 | 7.1 | 50 |
| 47_14 | 55.9 | 11.4 | 73 |
| 47_44 | 94.2 | 12.7 | 113 |
| 47_45 | 49.2 | 10.6 | 65 |
| 11_13 | 40.6 | 5.7 | 49 |
| 11_14 | 39.2 | 7.1 | 50 |
| 13_14 | 38.3 | 7.5 | 50 |
| 14_24 | 37.9 | 11.4 | 55 |
| 14_25 | 25.0 | 8.7 | 38 |
| 14_32 | 40.0 | 8.5 | 53 |
| 10_11 | 41.9 | 7.1 | 53 |
| 10_13 | 64.1 | 13.4 | 84 |
| 10_14 | 48.1 | 12.0 | 66 |
| 10_25 | 34.4 | 15.9 | 58 |
| 10_32 | 33.3 | 6.2 | 43 |
| 24_25 | 27.9 | 9.2 | 42 |
| 24_32 | 38.3 | 5.4 | 46 |
| 25_32 | 33.6 | 7.8 | 45 |

Chosen maximum lengths per tract after test runs described in *2.4 Tractography Parameters.* The first column corresponds to the regions the tract is reconstructed between. If a pair is not listed, there was no maximum length chosen after testing, as it either did not generate streamlines, or was not affected by varying parameters. M = mean; SD = standard deviation.
